# Targeting DNMT1 augments anti-tumor CD8⁺ T cell function

**DOI:** 10.64898/2026.07.03.736412

**Authors:** Maya G. Deshmukh, Danielle Sohai, Kamal Obbad, Koonam Park, Simon Milette, Peili Gu, Henna S. Nam, Andrew J. Daniels, Krasimir A. Spasov, Michael Hurwitz, Samuel Katz, Richard A. Flavell, Karen S. Anderson, Marcus Bosenberg, Goran Micevic

## Abstract

Chronic stimulation of CD8⁺ T cells within the tumor microenvironment (TME) induces a hypofunctional state characterized by diminished cytotoxicity and functionally impaired anti-tumor function, known as exhaustion. Exhaustion is associated with epigenetic changes that remain relatively stable despite interventions like immune checkpoint inhibition (ICI). Although epigenetic changes are potentially reversible, reports of therapeutic strategies to effectively restore function in exhausted CD8⁺ T cells remain limited. Here, we report DNA methyltransferase 1 (DNMT1) inhibition (DNMT1i) in counteracting CD8^+^ T cell dysfunction during the anti-tumor response. We show that DNMT1i synergizes with ICI to rescue the tumor cell killing activity of chronically stimulated CD8⁺ T cells in a melanoma model. DNMT1i mitigates transcriptional features of exhaustion while inducing a divergent effector program. DNMT1i attenuates the global increase in chromatin accessibility associated with exhaustion and enables epigenetic remodeling of the exhausted chromatin landscape upon restimulation. Finally, DNMT1i enhances the effector function of melanoma patient-derived tumor infiltrating lymphocytes after prolonged *ex vivo* expansion. These studies establish DNMT1 targeting as a promising strategy to counteract CD8⁺ T cell exhaustion and potentiate ICI efficacy.

## Introduction

The advent of ICI has transformed the therapeutic landscape of melanoma. Despite these advances, approximately half of patients with advanced melanoma treated with ICI do not derive long-term benefit (*1*). CD8⁺ T cell function underlies ICI responses, and T cell exhaustion in this compartment limits the responsiveness to ICI (*2, 3*). Our group and others have identified a subpopulation of functional memory anti-tumor CD8⁺ T cells that express interleukin-7 receptor (IL7R) that mediate superior tumor control in response to immune checkpoint blockade (*4, 5*).

IL7R^hi^ anti-tumor CD8⁺ T cells retain high expression of the transcription factor gene *Tcf7*, the ability to secrete effector cytokines, and, notably, a poised chromatin landscape with increased accessibility at the promoter of genes associated with cytotoxicity and memory relative to exhausted populations (*4*). These IL7R⁺ cells overlap functionally with progenitor exhausted (T_PEX_/TX_p_) CD8⁺ T cells, the self-renewing PD1⁺TCF1⁺TOX⁺ population that fuels the proliferative response to checkpoint blockade (*6*) However, progenitor frequency and effector output can be uncoupled, and strategies that expand progenitor-marked cells do not necessarily restore function, underscoring the need for interventions evaluated by functional rescue rather than marker frequency alone.

Epigenetic plasticity is a key feature of anti-tumor CD8⁺ T cells with functional potential. Chronic antigen stimulation and other tumor microenvironment (TME)-intrinsic cues initiate epigenetic changes in CD8⁺ T cells that precipitate terminal differentiation and preclude anti-tumor function in response to ICI (*7–9*). Selection and expansion of patient-specific IL7R^hi^ CD8⁺ T cells for adoptive cell therapy (ACT) is an attractive potential therapeutic strategy for melanoma, but IL7R^hi^ anti-tumor CD8⁺ T cells are rare relative to exhausted IL7R^lo^ anti-tumor CD8⁺ T cells (*4*). Given that *Tcf7* promoter accessibility is in part regulated by DNA methylation, we have previously found that treating IL7R^lo^ CD8⁺ T cells from tumor-draining lymph nodes with a DNA methyltransferase inhibitor can rescue their anti-tumor function (*4*). Among DNA methyltransferases, DNMT3A has been identified as a key driver of *de novo* methylation at stemness-associated loci, including the *Tcf7* promoter (*10*). DNMT3B, on the other hand, shows reduced expression in mature lymphocytes compared to DNMT3A and DNMT1 (*11*). The role of DNMT1, the maintenance methyltransferase responsible for propagating existing methylation marks during cell division, in CD8+ T cell exhaustion is yet unclear.

Here, we reveal a previously unappreciated role for DNMT1 in restraining anti-tumor function in chronically stimulated CD8⁺ T cells, which can be pharmacologically inhibited. We demonstrate that DNMT1i combined with ICI improves anti-tumor responses in a melanoma model and characterize the transcriptional and epigenetic changes driven by DNMT1 inhibition in chronically stimulated CD8⁺ T cells *in vitro*. Finally, patient-derived tumor-infiltrating lymphocytes (TILs) undergoing rapid expansion (REP) for melanoma ACT subjected to DNMT1i restore cytokine production and transcriptional features of cytotoxicity. Our findings delineate a novel role for DNMT1 in CD8⁺ T cell exhaustion in the context of chronic stimulation and propose DNMT1i as a strategy to maintain cytotoxic potential of anti-tumor CD8⁺ T cells in melanoma ICI.

## Results

### DNMT1 inhibition improves anti-tumor responses to ICI in a melanoma model

To evaluate the effect of combined DNMT1i and ICI therapy on the anti-tumor response, we used the YUMM-gp33 mouse melanoma cell lines (driven by clinically-relevant melanoma mutations *Braf*^V600E^*Cdkn2a*^-/-^) and immunocompetent C57Bl/6 mice (*12*). This model has demonstrated tumor outgrowth in immunocompetent mice that recapitulates progenitor exhausted (T_PEX_) and terminally exhausted T cell populations (T_EX_) seen in the human melanoma tumor microenvironment (TME) and responds to ICI (*4, 12*). In this model, dual ICI (anti-PD-1 and anti-CTLA-4) leads to 86% survival and anti-CTLA-4 alone leads to survival of about half of the treated mice at 60 days post injection (Fig S1A). We thus used ICI with anti-CTLA-4 for further experiments assessing adjuvant DNMT inhibitors.

We injected YUMM-gp33 cells subcutaneously into C57BL/6 mice. After allowing for the tumors to grow for 7 days, we treated the mice with anti-CTLA-4 (25 μg, ICI), RG108 (85 μg, DNMT1i) or combination of them (ICI+DNMT1i) by intraperitoneal injection every other day for four doses (Fig 1A). The DNMT1i group reached tumor volume endpoints by day 30, with no animals exhibiting tumor clearance (Fig 1B, left). ICI prolonged the overall survival compared to DNMT1i (Fig 1C, left). ICI+DNMT1i led to tumor clearance in seven out of nine mice (Fig 1B, right) and significantly longer survival compared to both DNMT1i and ICI (Fig 1C, left), demonstrating a superior anti-tumor response compared with ICI monotherapy.

**Figure 1.**
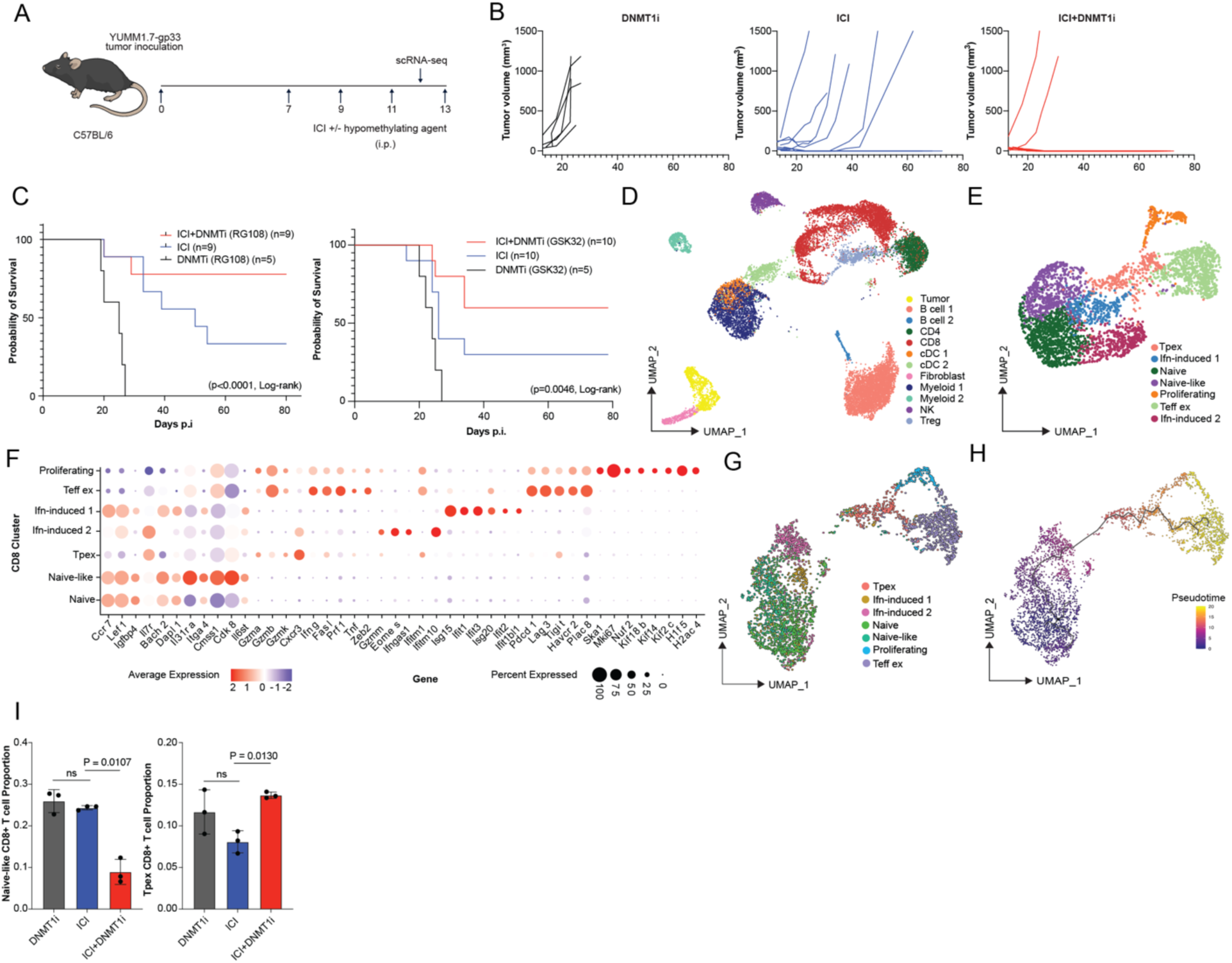
DNMT1i potentiates anti-CTLA4 therapy in a mouse model of melanoma. (A) Timeline for tumor injection and drug treatments. Tumor cells were injected on d0. (B) Tumor growth in individual mice treated with DNMTi (RG108) alone, ICI (anti-CTLA4) alone, or dual therapy (ICI+DNMTi). (C) Overall survival of mice receiving ICI (anti-CTLA4), DNMT1i (RG108 or GSK32), or ICI+DNMT1i (RG108 or GSK32) (log-rank Mantel Cox test). (D) UMAP reduction of identified clusters from 4 treatment groups (DNMTi alone (RG108), ICI (anti-CTLA4) alone, dual therapy, and control) with 3 mice per condition and 2 tissues (tdLN and tumor) from each mouse. (E) UMAP reduction of total CD8+ T cells from tdLN from all 4 treatment groups. (F) Dot plot showing expression of key markers defining each CD8 cluster. (G) UMAP reduction of total CD8+ T cells from tdLN from all 4 treatment groups using Monocle3 for trajectory analysis and (H) trajectory analysis colored by pseudotime. (I) Pseudobulk quantification of the proportion of CD8⁺ T cells in the naïve-like cluster (left) and T_PEX_ cluster (right) (n=3).

To confirm our observed phenotype was due to DNMT1 inhibition, we evaluated another DNMT1 selective inhibitor, GSK-3685032 (henceforth GSK32), and pan-DNMT inhibitors (SGI-1027, 5-Azacytidine, Decitabine) in combination with ICI in vivo. GSK32 has an IC_50_ of 0.036 μM and is about 2,500-fold more selective for DNMT1 than DNMT3A or DNMT3B (*13, 14*). We further validated that GSK32 specifically targets DNMT1 at the protein level. In agreement with a previous report showing that a GSK32 derivative targeted DNMT1 for degradation, we found that GSK32 treatment lead to reduced DNMT1, but not DNMT3a, in activated CD8^+^ T cells (Fig S1B). DNMT1i did not reduce DNMT1 in naïve cells, in which basal DNMT1 is lower at baseline (Fig S1B). This may be because DNMT1 is a maintenance methyltransferase and GSK-32 preferentially binds to DNMT1 when bound to hemimethylated CpG dyads. None of the tested pan-DNMT inhibitors improved the response to ICI (Figure S1C whereas combining GSK32 and ICI induced tumor clearance in 6/10 mice and significantly prolonged survival (*p* =0.0046, log-rank) (Fig 1C, right). These findings suggested that DNMT1i may enhance the effectiveness of ICI and suggested a role for DNMT1 in regulating immune cell function.

### DNMT1i in combination with ICI potentiates T_PEX_ formation in the tumor-draining lymph node

To investigate the effect of ICI+DNMT1i therapy on immune cell populations, we isolated immune cells from tumors and tumor draining lymph nodes (tdLNs) of mice treated with ICI, DNMT1i or ICI+DNMT1i and performed single-cell RNA sequencing (scRNA-seq) (Fig 1A). From 17,116 cells, we identified populations corresponding to B cells, myeloid cells, conventional dendritic cells (cDCs), NK cells, fibroblasts, and T cells in the tdLN (Fig 1D-F S1D). Given the central role of T cells in ICI response, we focused our analysis on T cells. Across 10 CD4⁺ T cell clusters identified in tdLNs and tumors, we did not detect significant differences in CD4⁺ cluster proportions between anti-CTLA4 and dual-therapy groups using a pseudobulk approach (Fig S1E). We next sub-clustered CD8⁺ T cells and identified 6 distinct CD8⁺ T cell populations: naïve-like, two interferon-induced populations, proliferating, progenitor exhausted (T_PEX_), and effector exhausted (T_EFF EX_) populations (*4, 15*), (Fig 1E,F).

To evaluate quantitative differences in the aforementioned CD8⁺ T cell clusters between the DNMT1i-treated, ICI-treated and dual-treated groups, we employed a pseudobulk approach to quantify the proportion of cells in each cluster. We found that the proportion of naïve-like CD8⁺ T cells was significantly reduced, approximately 2.7-fold (Welch’s *t*-test, *p* = 0.0107), in tdLNs of the ICI + RG108 group compared to those of the ICI group (Fig 1I, left). There was no significant difference in the proportion of naïve-like CD8⁺ T cells between the DNMT1i group and the ICI group, suggesting an effect specific to the combination of DNMT1i with ICI. We further found that the proportion of T_PEX_ CD8⁺ cells was significantly higher, approximately 1.7-fold (Welch’s *t*-test, *p* = 0.0130), in ICI+DNMT1i-treated mice than in ICI-treated mice (Fig 1I, right). We found no significant change in T_PEX_ proportion with DNMT1i monotherapy compared to ICI, again highlighting an effect specific to the combined immune-epigenetic (ICI+DNMT1i) therapy.

TdLN-resident T_PEX_ cells were defined by co-expression of *Tcf7* and *Pdcd1* together with the survival receptor *Il7r*, and were low for terminal-exhaustion marker *Havcr2*, consistent with consensus criteria for progenitor exhausted (T_PEX_/TX_p_) CD8⁺ T cells (*16, 17*). This population also expressed effector-associated transcripts (*Gzmb, Gzma, Gzmk, Cxcr3*), in keeping with the proliferation-competent, transitional nature of tdLN Tpex that seed effector progeny that can become progressively exhausted to T_EFF EX_ (*18, 19*).

(*3–5*). To investigate whether this trajectory was present in our dataset, we used pseudotime analysis of gene expression to order cells along a trajectory. Pseudotime analysis on tdLN CD8⁺ T cells yielded a trajectory starting with naïve CD8 T cells, progressing through a naïve-like state, interferon-induced 2, transitioning through T_PEX_ and terminating in T_EFF EX_ (Fig 1G,H). 3D pseudotime analysis further enabled visualization of the T_PEX_->T_EFF EX_ trajectory as progressing through a proliferating intermediate population (Fig S1F), consistent with previous reports that this progenitor population served as a proliferation-competent resource population that could terminally differentiate (*20, 21*). Ordering these populations along pseudotime thus validated the identification of a canonically immune checkpoint-responsive T_PEX_ population. The frequency of T_EFF EX_ in the tdLN and in the TME were not significantly impacted by dual therapy (Fig S1G).

Together, these findings demonstrate that DNMT1i+ICI is associated with enrichment in a T_PEX_-like population accompanied by reduction of an “upstream” naïve-like CD8⁺ T cell compartment. This suggests that dual DNMT1i+ICI therapy promotes the formation of T_PEX_ from naïve-like precursors.

### DNMT1 inhibition restores function of chronically stimulated CD8⁺ T cells

Our *in vivo* findings suggested that DNMT1 inhibition promoted the formation of T_PEX_ from naïve-like precursors during the ICI response, implicating DNMT1 in regulating anti-tumor CD8⁺ T cell differentiation. Terminal differentiation and dysfunction of CD8⁺ T cells in the anti-tumor response are associated with chronic exposure to antigens and other TME-specific factors (*22, 23*). To further evaluate the role of DNMT1i on CD8⁺ T cell function in the context of dysfunction-promoting chronic stimulation, we adapted a previously reported *in vitro* model of induction of dysfunction (*24*). Naïve, monoclonal gp33-reactive CD8⁺ T cells were isolated from P14 transgenic mice and stimulated with dendritic cells pulsed with gp33 peptide and cultured for one week with or without phorbol 12-myristate 13-acetate (PMA) and with or without DNMT1i (*24*) (Fig 2A). PMA activates protein kinase C (PKC) downstream of the T cell receptor, and chronic signaling following activation drives terminal differentiation and exhaustion (*24*). After one week of “rest” or chronic stimulation with PMA following initial priming with peptide-pulsed dendritic cells, CD8⁺ T cells were re-challenged with gp33 antigen to assess cytokine release or assessed for capacity to kill YUMM-gp33 cells. Henceforth, CD8⁺ T cells that were primed and rested for a week are denoted as memory-like or T_M_, whereas those chronically stimulated for a week following priming are denoted as T_EX_ (Fig 2B).

**Figure 2.**
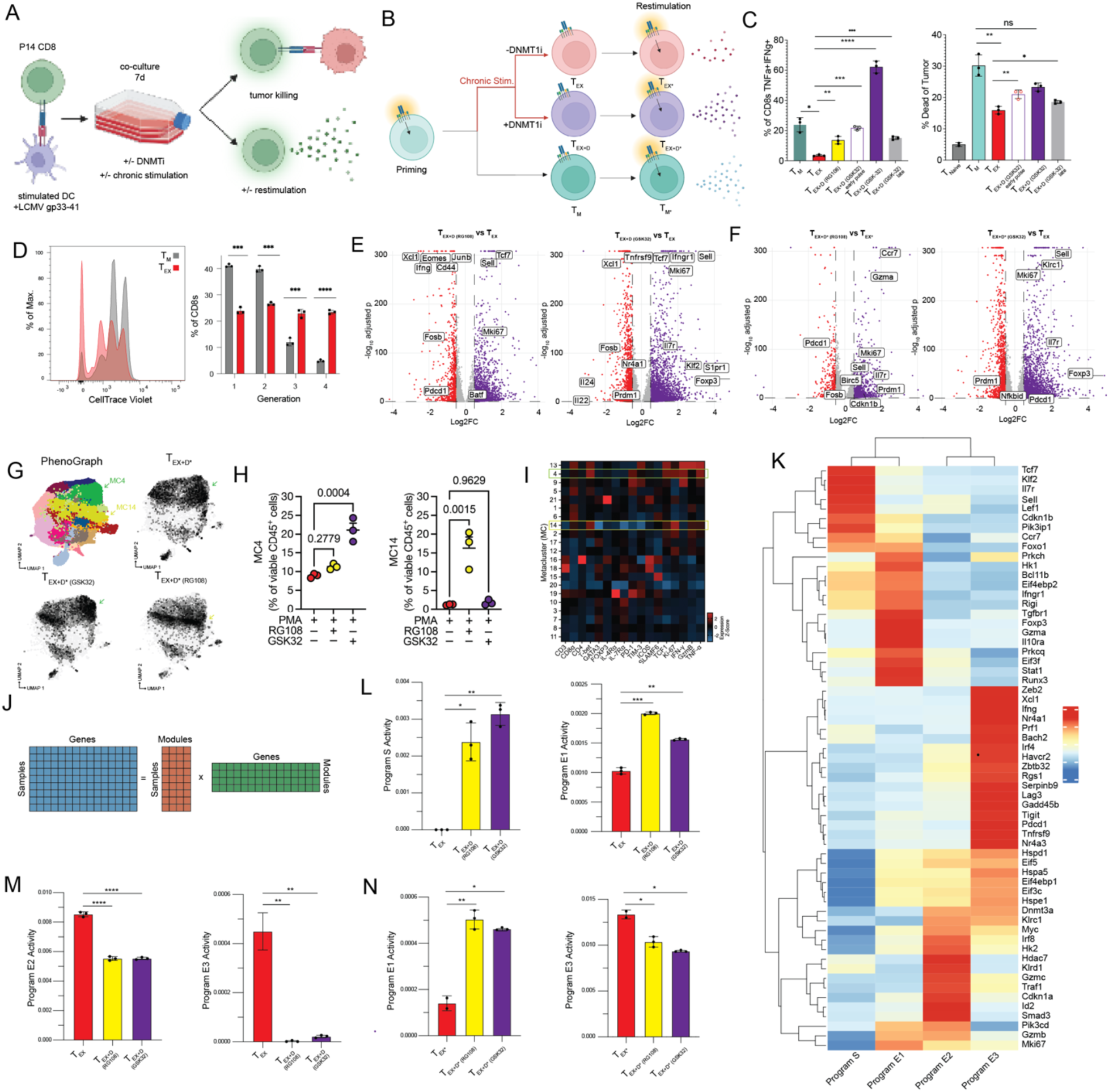
DNMT1i restores function of and transcriptionally reprograms chronically stimulated CD8+ T cells. (A) Schematic of *in vitro* primary P14 CD8+ T cell culture model. (B) Schematic describing abbreviated nomenclature for *in vitro* cell states (T_M_, T_EX_, T_EX+D_, T_M*_, T_EX*_, T_EX+D*_). (C) P14 CD8⁺ T cells were isolated on d7 of culture and restimulated with gp33 peptide for flow cytometry analysis of IFNγ and TNFα production (left) or co-cultured with gp33-expressing mouse melanoma line YUMM-G1-gp33 to assess tumor cell killing (right) by flow cytometry (n=3). (D) Chronic stimulation promotes proliferation of CD8+ T cells as shown by CellTrace Violet dilution (left) and quantified as proportion of cells in each generation (right) (n=3). (E-F) Violin plots showing differentially expressed genes between T_EX+D_ and T_EX_ (E) and between T_EX+D*_ and T_EX*_ (F) with RG108 (left) or with GSK32 (right). An adjusted p-value cutoff of 0.05 and log2FC cutoff of 0.5 were used. (G) UMAP depicting clusters obtained using PhenoGraph for concatenated samples (top left) and density plots for TEX* (top right), TEX+D* (GSK-32) (bottom left), and TEX+D* (RG108). (H) Proportion of live, CD45+ cells in MC4 (left) and MC14 (right) across samples. (I) Heatmap representing the relative expression of markers across clusters. (J) Schematic showing consensus nonnegative matrix factorization (cNMF) matrices. (K) Gene by program matrix from cNMF showing loadings for genes in the top 150 ranked gene list for each module. (L) Comparison of T_EX_ vs T_EX+D_ usage of Program S (left) and Program E1(right). (M) Comparison of T_EX_ vs T_EX+D_ usage of Program E2 (left) and Program E3 (right). (N) Comparison of T_EX*_ vs T_EX+D*_ usage of Program E1 (left) and Program E3 (right).

Chronic stimulation led to a 6.7-fold reduction in the percentage of TNFα⁺IFNγ⁺ CD8⁺ T cells upon re-challenge (T_M_ vs T_EX_, *p* = 0.0161) (Fig 2C. Chronic stimulation after priming also attenuated cytotoxicity, leading to a 1.9-fold reduction in killing (T_M_ vs T_EX_, *p* = 0.0024) (Fig 2C. Despite functional impairment, chronic stimulation increased cell proliferation (Fig 2D), in line with previous reports that effector function and expansion are uncoupled in dysfunctional CD8⁺ T cells (*25*).

We next evaluated the ability of DNMT1i to prevent or rescue chronic stimulation-induced dysfunction. PMA and DNMT1i-treated CD8⁺ T cells are referred to as T_EX+D_ and re-challenged CD8⁺ T cells are denoted with an asterisk (T_M*_, T_EX*_, T_EX+D*_) (Fig 2B). To ask whether DNMT1i could prevent functional exhaustion, we continuously treated CD8⁺ T cells undergoing PMA-induced chronic stimulation with RG108 or GSK32. GSK32 increased the percentage of TNFα⁺IFNγ⁺ CD8⁺ T cells 17-fold (*p* < 0.0001), fully restoring this population, and increased the killing capacity (1.5-fold increase, *p* = 0.0019), RG108 also significantly, although less effectively, increased TNFα⁺IFNγ⁺ CD8⁺ T cells but it had no effect on cytotoxicity (Fig. 2C, Fig S2B).

It was recently shown that both knockout of *DNMT1* and pharmacologic inhibition with a GSK32 analog in human T cells leads to NK-like properties with enhanced cytotoxic function (*26*). To determine whether DNMT1i was sufficient to increase cytotoxicity, we treated primed CD8⁺ T cells continuously with GSK32 in the absence of chronic stimulation for a week before challenging with tumor cells and found that DNMT1i treatment did not enhance the cytotoxicity (Fig S2C). In this setting, DNMT1 appears to play a role in regulating the response to chronic stimulation that may be uncoupled from its lineage-defining role.

*In vitro* experiments with DNMT1i thus far have involved continuously treating with DNMT1i concurrently with chronic stimulation. We next tested the ability of DNMT1i to attenuate functional exhaustion when provided prior to or after chronic stimulation. Anti-tumor CD8⁺ T cells can acquire dysfunction-associated epigenetic programs as early as 6 hours following activation, and these changes are reinforced progressively over days (*25*). To determine if early DNMT1i could attenuate features of exhaustion during chronic stimulation, we performed an early, transient pulse with GSK32 during the first 36 hours of stimulation. This one-time early pulse of DNMT1i restored the frequency of TNFα⁺IFNγ⁺ CD8⁺ T cells and sustained tumor killing capacity despite continued PMA stimulation following the DNMT1i pulse (Fig 2C). Early DNMT1i was thus capable of maintaining cytotoxic potential and preserving functional tumor cell killing despite continued stimulation and thus counteracts features of exhaustion. Conversely, when CD8⁺ T cells were chronically stimulated without DNMT1i for one week, DNMT1i at the time of rechallenge was sufficient to increase the frequency of TNFα⁺IFNγ⁺ CD8⁺ T cells and tumor cell killing (Fig 2C,D. This suggests that DNMT1i may reverse features of functional exhaustion even after CD8⁺ T cells have received prolonged chronic stimulation cues.

### DNMT1i induces a stem-like transcriptional program in chronically stimulated CD8+ T cells

To evaluate transcriptional changes by which DNMT1i counteracted early dysfunction, we performed bulk RNA sequencing on CD8⁺ T cells from the *in vitro* chronic stimulation model (Fig 2A). Principal component analysis (PCA) revealed that samples clustered primarily according to activation and exhaustion state, though DNMT1i shifted the transcriptome of chronically stimulated CD8⁺ T cells (T_EX+D_) towards the T_M_ cluster along both PC1 and PC2 (Fig S2D).

We first analyzed transcriptional changes induced by chronic stimulation alone (T_EX_ vs T_M_, 8,589 differentially expressed genes). Chronic stimulation increases transcription of exhaustion-and effector-associated genes (*Ifng, Hk2, Havcr2, Prdm1, Dnmt3a*) and downregulated genes associated with quiescence (*Tcf7*, *Klf2*), cytokine responsiveness (*Xcl1, Ifngr1*) and early effector differentiation (*Gzmk, Gzma*) (S2E,G). Gene set enrichment analysis (GSEA) on T_M_ vs T_EX_ further revealed that chronic stimulation enriched Myc targets and TNFα signaling via NFKB while downregulated IFNγ and IFNα response pathways (Fig S2F), suggesting polarization of effector function with chronic stimulation.

We next compared T_EX_ to T_EX+D_ (7,836 DEG with RG108; 8,661 DEG with GSK32). T_EX+D_ treated with either RG108 or GSK32 upregulated *Mki67*, *Tcf7* and *Sell* while downregulating activation-associated *Fosb* and *Xcl1* relative to T_EX_ (Fig 2E. RG108 treatment downregulated activation-associated (*Junb*, *Pdcd1*, *Cd44*) and effector-associated (*Ifng*, *Eomes*) genes (Fig 2E, left). GSK32 treatment uniquely differentially upregulated specific quiescence-associated genes (*Il7r, Klf2*) and downregulated a unique set of activation- and exhaustion-associated genes (*Tnfrsf9*, *Nr4a1*, *Prdm1*) (Fig 2E, right). GSK32 treatment also upregulated *Foxp3*, which is a known target of DNMT1 (*27*). Taken together, this suggested that DNMT1i dampened activation- and effector-associated gene programs while enforcing quiescence-associated gene programs in the setting of chronic stimulation. Restimulation of chronically stimulated CD8⁺ T cells induced 8,643 DEG (T_EX_ vs T_EX*_). In this context, RG108 treatment induced 8,095 DEG and GSK32 induced 8,121 DEG. Upon restimulation, DNMT1i enabled sustained expression of *Il7r*, *Sell* and *Mki67* in chronically stimulated CD8⁺ T cells (Fig 2F). We assessed the impact of DNMT1i-induced transcriptional changes at the level of expression using high-dimensional spectral flow cytometry (Fig 2G). In T_EX+D*,_ GSK32 induced enrichment of meta-cluster 4, marked by high expression of T-bet, PD-1, TNFα, IFNγ, Ki-67, and TCF1 (Fig 2H,I, S2J). RG108 induced enrichment of meta-cluster 14, marked by high T-bet, Ki-67, GzmB, TNFα, and TCF1 expression (Fig 2H,I, S2K). Together, these data further suggested that DNMT1i preserved an early activation phenotype on restimulation in a subset of chronically stimulated CD8⁺ T cells.

To characterize differences in effector programs underlying DNMT1i-mediated functional restoration of chronically stimulated CD8⁺ T cells, we used consensus non-negative matrix factorization (cNMF). cNMF trains latent gene programs defined by sets of co-expressed genes and their relative usage across samples (Fig 2J) (*28*). We identified one stemness-associated (Program S) and three effector-associated (E1, E2, E3) programs. Program S was characterized by quiescence-associated gene loadings (*Tcf7, Klf2, Il7r, Sell, Lef1, Ccr7, Foxo1, Pik3i1p1*) (Fig 2K). In the hierarchal clustering dendrogram, Program E1 clustered most closely to Program S, with associated genes related to stemness (*Ccr7, Foxo1*), interferon-responsiveness (*Rigi*, *Stat1, Ifngr1*), early effector function (*Runx3*, *Gzma, Gzmb*) and biosynthetic regulation (*Eif4ebp2, Eif3f*) (Fig 2K). Programs E2 and E3 clustered together, with E2 characterized by genes associated with NK-like function (*Klrc1*, *Klrd1*) and effector differentiation and exhaustion (*Hk2*, *Pik3cd*, *Gzmb, Id2*) (Fig 2K). E3 was further associated with genes characterizing activation and exhaustion (*Xcl1, Ifng, Prf1, Serpinb9, Havcr2, Lag3, Tigit, Pdcd1*) and biosynthetic activity (*Eif5, Eif4ebp1, Eif3c, Hspd1, Hspe1*) (Fig 2K). E1 thus most closely resembled an early effector-memory program, while E2 and E3 represented effector gene programs associated with terminal differentiation.

Evaluation of the usage of each program across conditions revealed that T_EX+D_ preferentially used Program S and E1 whereas T_EX_ preferentially used E2 and E3, supporting the role of DNMT1i in restraining effector- and activation-associated transcriptional programs under chronic stimulation signals (Fig 2L,M). Upon restimulation, DNMT1i-treated CD8⁺ T cells show higher usage of E1 and a modest decrease in E3 usage (Fig 2N).

Next, we sought to validate whether induction of divergent effector programs by DNMT1i during chronic stimulation is due to on-target DNMT1 inhibition and whether DNMT1i in mouse cells induces transcriptional changes that overlap with DNMT1 knockout in human T cells. We compared differentially expressed genes by pharmacologic DNMT1i from our mouse data with DNMT1 knockout in human T cells from Li et al. (*26*) (Fig S2H). We compared differentially expressed genes induced by DNMT1i in memory-like (T_M_ vs T_M+D_), exhausted (T_EX_ vs T_EX+D_), and exhausted-restimulated (T_EX*_ vs T_EX+D*_) to differentially expressed genes in human DNMT1 KO (hDNMT1 KO vs WT) from Li et al. and assessed overlap of DEG(*26*). DNMT1KO in human T cells and DNMT1i in mouse T cells regulate similar sets of genes, including shared upregulation of *Gzma, Prf1, Nfatc3* and *Nkg7* and downregulation of *Myc*, *Nfatc1*, *Rela, Hk2*, and *Atf4*. (Fig S2I).

Together, these data suggested that DNMT1 was involved in patterning early fate choices in effector CD8⁺ T cells under chronic stimulation. DNMT1i insulated T_EX+D_ from transcription of effector-associated modules (E2 and E3) under chronic stimulation. Upon restimulation, DNMT1i-treated CD8⁺ T cells retained superior effector function associated with use of a divergent transcriptional effector program (E1) characterized by reduced biosynthetic and terminal effector transcriptional activity compared with E3, which was more strongly induced in T_EX_.

### DNMT1i increases epigenetic plasticity of chronically stimulated CD8⁺ T cells

We next asked 1) whether *in vitro* T_EX_ were epigenetically ‘scarred’ relative to T_M_ and, if so, 2) whether DNMT1 inhibition affects the epigenetic landscape using assay for transposase-accessible chromatin with high-throughput sequencing (ATAC-seq) (*29*).

Differential chromatin accessibility was assessed across a consensus set of 118,684 peaks. T_EX_ had distinct chromatin accessibility relative to T_M_ (71,770 differentially accessible peaks) in regions affecting genes associated with exhaustion and quiescence (*Havcr2*, *Tox2*, *Sell, Runx3, Id2*). Globally, differentially regulated peaks became, on average, more accessible with exhaustion, in agreement with prior reports (Fig 3A, S3B)(*30*). We next compared the epigenetic landscape of T_EX+D_ with T_EX_. Principal component analysis showed that T_M_ clustered separately from T_EX_ and T_EX+D_, suggesting that DNMT1i resulted in overall chromatin accessibility pattern closer to T_EX_ than T_M_ (Fig 3B). T_EX+D_ differentially regulated 14,706 peaks and was associated with an overall global decrease in chromatin accessibility relative to T_EX_ (Fig 3A). Of these, 96 significantly gained accessibility with DNMT1i (Fig 3C). T_EX+D_ lost accessibility at 14,610 peaks relative to T_EX_, the majority (76%) overlapped with chromatin accessibility lost in T_M_ relative to T_EX_ (Fig 3C). Shared accessibility patterns between T_M_ and T_EX+D_ were mapped to genes involved in T cell differentiation (*Runx3, Egr3, Ccr7, Tcf7, Tbx21*) by overrepresentation analysis (Fig S3A). T_M_ showed reduced accessibility at 5-fold more peaks relative to T_EX_ that did not overlap with those lost by T_EX+D_, overrepresented for processes involved in phosphorylation, signal transduction and differentiation (Fig 3C). Thus, despite focal re-patterning and some shared regulation of T_EX_-induced peaks by T_EX+D_ and T_M_, T_EX+D_ appear globally similar to T_EX_ in terms of chromatin accessibility (Fig 3D).

**Figure 3.**
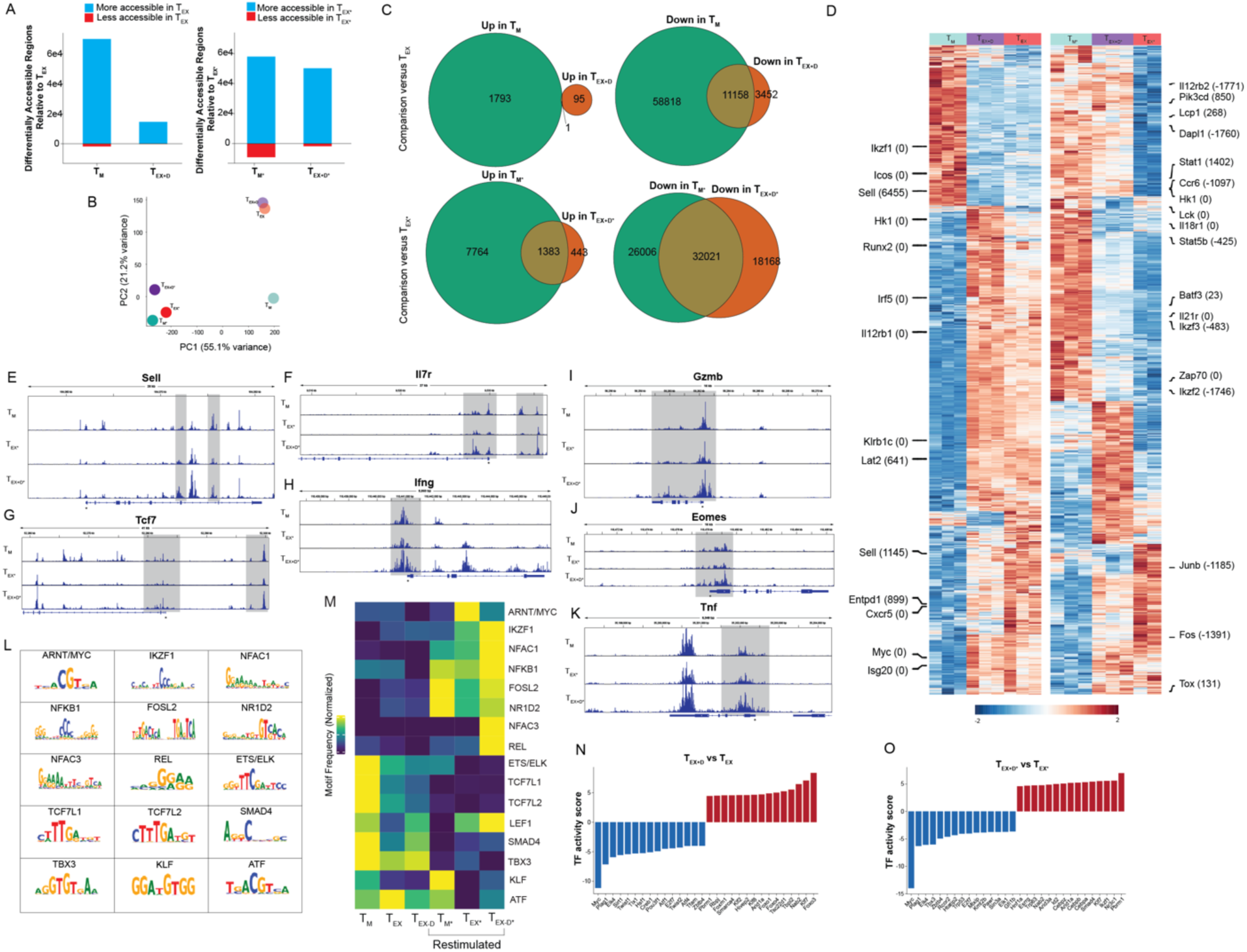
DNMT1i modulates the epigenetic and regulatory landscape of chronically stimulated CD8⁺ T cells *in vitro*. (A) Barplot showing number of total peaks differentially accessible between T_M_ and T_EX_ (left plot, left bar), T_EX+D_ and T_EX_ (left plot, right bar), T_M*_ and T_EX*_ (right plot, left bar), and T_EX+D*_ and T_EX*_ (right plot, right bar). (B) PCA of T_M_, T_EX+D_, T_EX_, T_M*_, T_EX+D*_, and T_EX*_ based on chromatin accessibility. (C) Venn diagrams showing overlap of genes upregulated in T_M_ and T_EX+D_ versus T_EX_ (top left), downregulated in T_M_ and T_EX+D_ versus T_EX_ (top right), upregulated in T_M*_ and T_EX+D*_ versus T_EX*_ (bottom left), and downregulated in T_M*_ and T_EX+D*_ versus T_EX*_ (bottom right). (D) Heatmap of peaks differentially regulated between T_M_, T_EX+D_, and T_EX_ (left) and between T_M*_, T_EX+D*_, and T_EX*_ (right) with select peaks annotated to immune-related gene regulatory regions and distance to TSS labeled (adj p-val < 0.05). (E-K) Genome browser tracks for T_M_, T_EX+D*_, and T_EX*_ for loci near *Sell* (E), *Il7r* (F), *Tcf7* (G), *Ifng* (H), *Gzmb* (I), *Eomes* (J) and *Tnf* (K) with TSS, gene boundaries and direction denoted below each plot. (L) Transcription factor motif families learned by ChromBPNet. (M) Heatmap showing frequency of learned motifs near genes across samples. (N-O) Decoupler regulon analysis of bulk RNA-seq profiles of T_EX+D_ vs T_EX_ (N) and T_EX+D*_ vs T_EX*_ (O). Gene names on x-axis denote TF regulons.

The chromatin accessibility patterns in T_M*_, T_EX*_ and T_EX+D*_ changed profoundly upon restimulation (Fig 3D). Restimulation decreased the Euclidean distances between T_EX*_, T_EX+D*_ and T_M*_, and DNMT1i induced a shift from the exhausted towards a memory-like state in PCA space (Fig 3B). Like T_M_ vs. T_EX_, T_M*_ showed a more closed chromatin landscape compared to T_EX*,_ with 86% of peaks less accessible (Fig 3A). T_EX+D*_ again primarily induced loss of accessibility relative to T_EX*_, with 64% of closed peaks shared with T_M_* (Fig 3A,C). This shared set mapped to genes overrepresented for differentiation (*Irf4*, *Ccr7, Runx2, Icos, Il7r*), phosphorylation, and signal transduction terms (Fig S3C).

Notably, T_EX+D*_ showed increased chromatin accessibility at key regulatory elements associated with CD8⁺ T cell phenotype. For example, an intronic and exon-containing region within the *Sell* locus regained accessibility in T_EX+D*_ compared to T_EX*_ (Fig 3E). Further, a distal regulatory region and TSS-containing region of *Il7r* regained accessibility in T_EX+D*_ relative to T_EX*_ (Fig 3F). A distal regulatory region and the proximal promoter and TSS of *Tcf7* retained accessibility in T_EX+D*_ relative to T_EX*_ (Fig 3G). DNMT1i similarly increased the accessibility in the proximal promoter for differentiation-associated transcription factor gene *Eomes* upon restimulation (Fig 3J). A similar pattern was seen for effector-associated genes *Gzmb*, *Ifng*, and *Tnf*. (Fig 3H,I,K). Together, these data indicate that while DNMT1i induces only modest and focal chromatin remodeling at rest, it enables widespread accessibility changes at memory- and effector-associated loci upon restimulation, consistent with a function in potentiating epigenetic plasticity in chronically stimulated CD8+ T cells.

### DNMT1i regulates TF activity of CD8⁺ T cells under chronic stimulation

Our data showed that DNMT1 inhibition in exhausted T cells induced focal re-patterning in chromatin accessibility of key regulatory loci (concentrated at memory- and exhaustion-associated loci) that likely drove global transcriptional remodeling and strong functional rescue. To characterize transcription factor (TF) families involved in regulatory changes induced by DNMT1i, we trained ChromBPNet models, a deep learning architecture for motif footprinting, and determined the frequency of learned motifs in peaks near genes (*31*) (Fig 3L). As expected, TCF7 motif families occurred more frequently near genes in the memory condition and activation-associated TF motifs (NFAC1, NFKB1, FOSL2, NR1F2) occurred more frequently in restimulated samples (Fig 3M).

At rest, T_EX_ showed reduced frequency of quiescence-associated motif families (KLF, LEF1, TCF7, and ETS/ELK) and increased frequency of FOSL2, MYC, and ATF families relative to T_M_ (Fig 3M. Upon restimulation, T_EX_ showed lower frequency of activation-associated (FOSL2, NR1D2, NFKB1) and quiescence-associated (ETS/ELK, KLF) families and increased frequency of MYC (Fig 3M). At rest, DNMT1i increased frequency of LEF1 motifs, but notably did not impact TCF7 family motif frequency (Fig 3M). Regulon analysis further identified significant upregulation of targets of Forkhead (FOX) and Kruppel-like factor (KLF) families, including quiescence-associated TFs Klf2 and Foxo1, with DNMT1i (Fig3N). On restimulation, DNMT1i partially restored frequencies of activation-associated motifs lost in T_EX_ relative to T_M_ (FOSL2, NFKB1, NR1D2) as well as KLF motifs (Fig 3N). Both at rest and on restimulation, DNMT1i decreased frequencies of MYC motifs and increased frequencies of IKZF1 and KLF motifs (Fig 3N). Regulon analysis confirmed significant transcriptional upregulation of the Ikzf1, Foxo1/3, and Klf2/7/8 regulons and downregulation of the Myc regulon with DNMT1i (Fig 3O).

Thus, despite limited global chromatin changes in T_EX+D_ relative to T_EX_, DNMT1i was associated with different TF activity under chronic stimulation. Footprinting and regulon analysis identified that DNMT1 drove differential activity of Myc, Atf, Klf, Lef1, and Ikzf1 TF motif families.

### DNMT1i drives global hypomethylation in CD8⁺ T cells

We next assessed methylation at the single CpG-site level across more than 285,000 sites in T_M_, T_EX_, and T_EX+D_ to determine whether DNMT1i-induced changes to regulatory TF activity were associated with DNA hypomethylation. In contrast to PCA of ATAC-seq, CpG methylation samples clustered according to drug treatment rather than exhaustion history (Fig 4B). Notably, there were no differentially regulated CpG sites between T_EX_ and T_M_ (S4A). DNMT1i-treated samples demonstrated global hypomethylation in a dose-dependent manner (Fig 4A). T_EX+D_ and T_M+D_ treated with an early pulse of GSK32 exhibited reduced methylation relative to T_EX_ and T_M_, validating that early demethylation can dictate long-term changes after the drug has been removed (Fig 4A). DNMT1i continuously treated T_EX_ and T_M_ demonstrated profound hypomethylation, including within regulatory regions of genes associated with CD8 quiescence (*Sell, Lef1, Il7r, Klf6, Foxo1, Bach2*) (Fig 4C,S4C-H).

**Figure 4.**
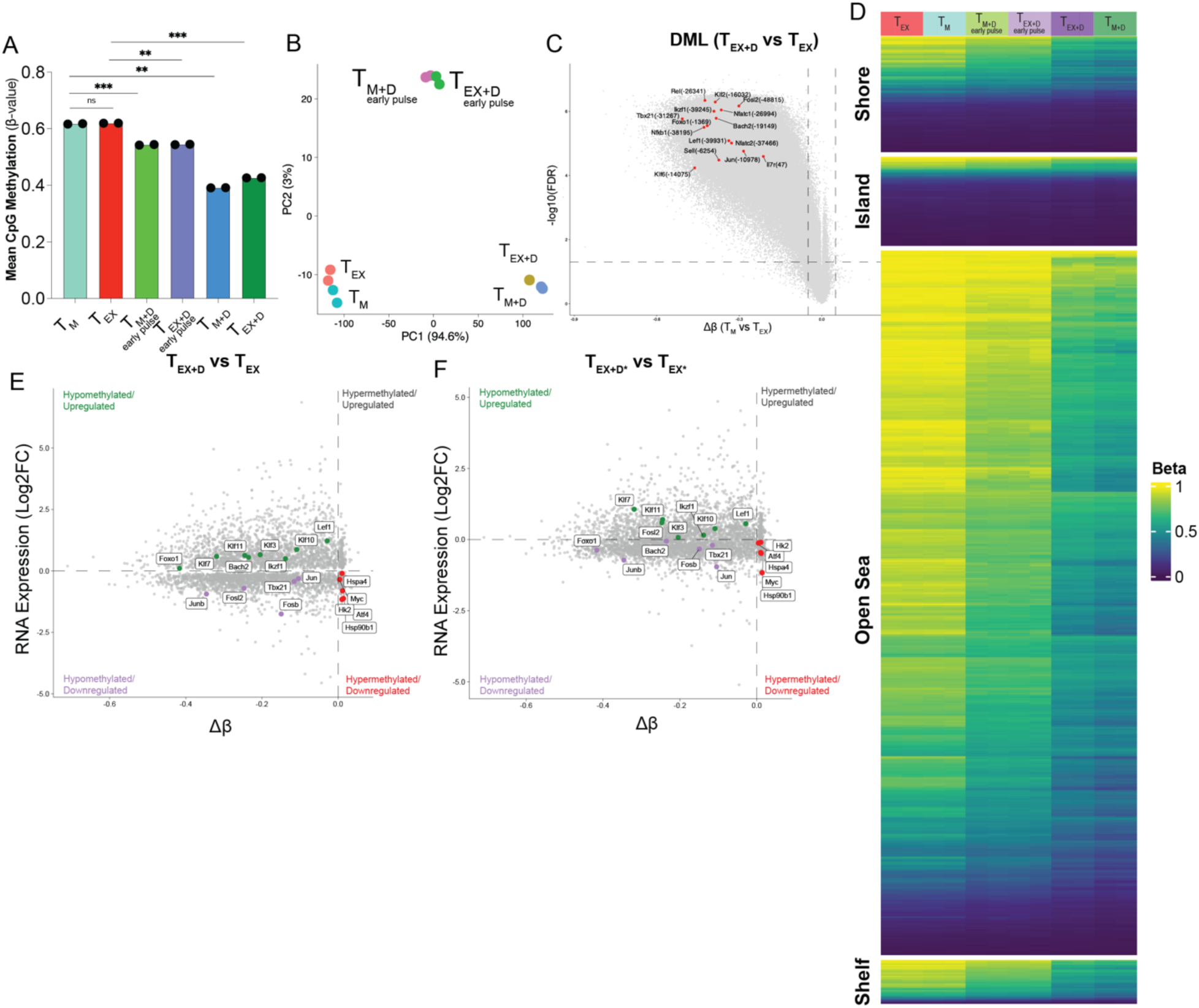
DNMT1i drives global DNA hypomethylation in chronically stimulated CD8^+^ T cells. (A) Mean global CpG methylation (*β* value) across all loci across treatment groups. (B) PCA of treatment groups based on methylation across all loci (M value). (C) Volcano plot of differentially methylated loci between T_EX+D_ vs T_EX_. (D) Heatmap showing *β* values for loci organized by CpG context across treatment groups. (E-F) Concordance plots comparing methylation and RNA seq for T_EX+D_ vs T_EX_ (left) and T_EX+D*_ vs T_EX*_ (right). Each locus within a CpG shelf, shore, or island and within −5kb to +1kb from a TSS was mapped to a gene. Methylation at that site relative to gene expression was plotted and defined by quadrant.

The majority of differentially methylated loci (DML) were located in open sea regions (76%) while fewer were located in CpG shores, shelves and islands (13%, 7%, and 4%, respectively) (Fig 4D, S4B). We next sought to determine whether site-specific changes in methylation were associated with changes in gene transcription. To constrain CpGs to those that were likely to regulate proximal genes, we included only DML mapped to island, shelf, or shore regions that were also within 5kb upstream to 1kb downstream from TSS of a given gene. As expected, most of these DML were hypomethylated both at rest and on restimulation (Fig 4E,F). Despite global hypomethylation, there were similar numbers of transcriptional upregulated and downregulated genes, reflecting a complex regulatory environment (Fig 4E,F). To determine which DNA methylation changes were more likely to have a functional effect, we integrated DNA methylation changes with gene expression (RNA-seq) data for each CD8 state. Using this integrated dataset, we found that *Lef1*, *Ikzf1*, and members of the *Klf* (*Klf3,7,10,11*) and *Fox* (*FoxO3,4*) were hypomethylated and upregulated in T_EX+D_ versus T_EX_ and T_EX+D*_ versus T_EX*_ (Fig 4E,F). Conversely, *Myc* and *Atf4* were consistently hypermethylated and downregulated (Fig 4E,F). These findings supported footprinting and regulon data nominating these TFs as potential key players in regulating the DNMT1i-induced state in chronically stimulated CD8⁺ T cells.

### DNMT1i de-represses *Il7r* and potentiates IL-7R–STAT5 signaling

Among the quiescence-associated loci de-repressed by DNMT1i, we focused on *Il7r*, given that IL-7R marks anti-tumor memory CD8⁺ T cells with superior efficacy (4) and is epigenetically regulated. We therefore integrated our transcriptional, methylation, and chromatin data at the *Il7r* locus and functionally interrogated its downstream signaling pathway.

DNMT1i increased *Il7r* transcription specifically when applied to CD8^+^ T cells in the exhausted state (Fig 5A). Relative to T_EX_, both RG108 and GSK32 upregulated *Il7r* at rest (T_EX+D_) and upon restimulation (T_EX+D*)_ (Fig. 5A). Notably, DNMT1i did not increase *Il7r* in memory-like cells (T_M_) and GSK32 modestly reduced it (T_M+D_), consistent with *Il7r* already being expressed in T_M_ and with DNMT1i-induced *Il7r* upregulation unique to the exhausted state (Fig. 5A).

**Figure 5.**
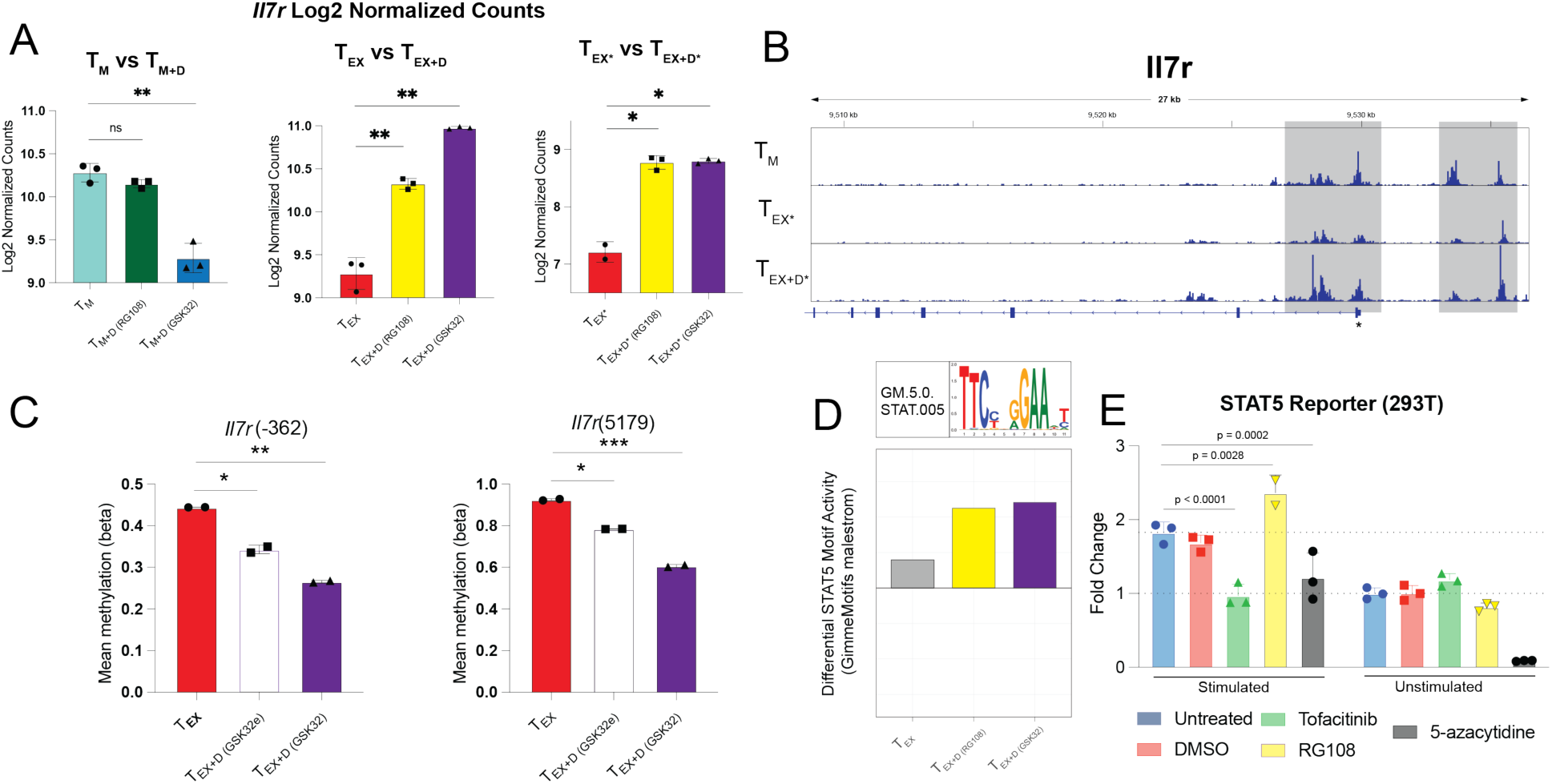
DNMT1i de-represses *Il7r* and potentiates IL-7R–STAT5 signaling in exhausted CD8⁺ T cells. (A) *Il7r* log2-normalized counts (bulk RNA-seq) for T_M_ vs T_M+D_, T_EX_ vs T_EX+D_, and T_EX*_ vs T_EX+D*_ with RG108 and GSK32 (n=3; adjusted p-values shown). (B) Mean DNA methylation (β value) at *Il7r* regulatory CpGs [*Il7r*(−362), promoter-proximal; *Il7r*(5179), intragenic] in T_EX_, T_EX+D_ (early GSK32 pulse, "GSK32e"), and T_EX+D_ (continuous GSK32). (C) Genome browser tracks of chromatin accessibility at the *Il7r* locus for T_M_, T_EX*_, and T_EX+D*_, with distal regulatory and TSS-proximal regions highlighted. (D) Differential STAT5 motif activity (GimmeMotifs maelstrom) in T_M_, T_EX*_, and T_EX+D*_ (RG108) and T_EX+D*_ (GSK32)); STAT5 motif logo shown above. (E) STAT5 reporter activity in 293T cells under IL-7 stimulation or unstimulated conditions, treated with vehicle (untreated), DMSO, tofacitinib (JAK/STATi), RG108 (DNMT1i) or 5-azacytidine (n=3; p-values vs untreated). *Statistics/tests as indicated*.

This de-repression was accompanied by focal epigenetic remodeling of the *Il7r* locus. DNMT1i reduced DNA methylation at *Il7r* regulatory CpGs, including a proximal promoter site [*Il7r*(−362)] and an intragenic site [*Il7r*(5179)], in a regimen-dependent manner, with continuous GSK32 producing greater hypomethylation than an early GSK32 pulse (Fig. 5C). In parallel, distal regulatory and TSS-proximal regions of *Il7r* regained chromatin accessibility in restimulated DNMT1i-treated cells (T_EX+D*)_ relative to T_EX_ (Fig. 5D), mirroring the genome-wide, restimulation-dependent re-opening described above.

To connect this locus-level remodeling to IL-7R signaling, we examined STAT5, the principal transcription factor downstream of IL-7R and a recognized regulator of exhausted CD8⁺ T cell differentiation (*6*). Differential motif analysis of our ATAC-seq data showed increased STAT5 motif activity in DNMT1i-treated exhausted cells (T_EX+D_, RG108 and GSK32) relative to T_EX_ (Fig. 5D). To test whether DNMT1i functionally augments STAT5-dependent transcription, we used a STAT5 reporter (293T). Under stimulation, RG108 significantly increased STAT5 reporter activity relative to untreated and vehicle controls (p = 0.0028), whereas the JAK inhibitor tofacitinib abolished the signal (p < 0.0001), confirming JAK/STAT dependence (Fig. 5E). The nucleoside analog 5-azacytidine did not potentiate, and significantly reduced, reporter activity (p = 0.0002), with near-complete loss of signal in unstimulated cells consistent with its cytotoxicity and with our finding that nucleoside pan-DNMT inhibitors lack the beneficial DNMT1i phenotype (Fig. S1C). Potentiation required stimulation, as no treatment increased STAT5 activity in unstimulated cells (Fig. 5E).

Together, these data identify *Il7r* as a locus of DNMT1i-driven hypomethylation, re-expression, and chromatin re-opening in exhausted CD8⁺ T cells, and show that DNMT1i potentiates stimulation-induced STAT5 activity and supporting a model in which DNMT1 inhibition restores competence of the IL-7R–STAT5 axis otherwise repressed under chronic stimulation.

### DNMT1i enhances effector function of patient-derived TILs from metastatic melanoma

Our findings implicated DNMT1 in enforcing exhaustion-associated programs early in chronic stimulation of CD8⁺ T cells and showed that DNMT1i attenuated dysfunction *in vivo* and *in vitro*. Further, we showed that DNMT1i can rescue cytokine production and cytotoxicity even when provided after chronic stimulation *in vitro*, suggesting that this approach may be useful for the functional reinvigoration of exhausted tumor infiltrating lymphocytes (TILs). Epigenetic reprogramming of patient-derived adoptive cell therapy (ACT) products is an attractive clinical strategy as it circumvents the potential challenges associated with systemic DNMT1i therapy, for which the therapeutic index in tumor-specific CD8⁺ T cells is likely unfavorable. ACT with patient-derived tumor infiltrating lymphocytes (TILs) is approved for patients with stage III/IV melanoma that have progressed on prior systemic therapy(*32*). While TILs are abundant and routinely isolated, tumor-reactive TILs are highly differentiated following the rapid expansion protocol (REP) used to expand the population for therapy. We thus sought to evaluate whether DNMT1i could improve the function of patient-derived TILs expanded *ex vivo*.

We expanded TILs from 5 patients with metastatic melanoma (Table S1) who had progressed on anti-PD-1 or anti-PD-L1 therapy alone or in combination with anti-CTLA4 in two phases: 1) initial expansion or “C1” and 2) rapid expansion with irradiated feeder cells, anti-CD3/anti-CD28 stimulation and IL2 or “C2” (Fig 6A) (*33*). We then further expanded C2 TILs (“post-C2”) in the presence or absence of GSK32 to determine whether DNMT1 inhibition during late-stage stimulation and expansion of highly differentiated TILs could restore effector function. Since patient 1 TILs were predominantly CD4 (75.4%), they were excluded from functional and transcriptional analysis (Fig S6A). We collected continuously stimulated post-C2 TILs from patients 2, 4, 5, and 6 and re-stimulated them (Fig 6B, C). Treatment of all four patient TILs with GSK32 enhanced the proportion of TNFα⁺IFNγ⁺ CD8⁺ T cells upon restimulation (*p* = 0.03). DNMT1i could thus restore features of effector function in patient-derived TILs.

**Figure 6.**
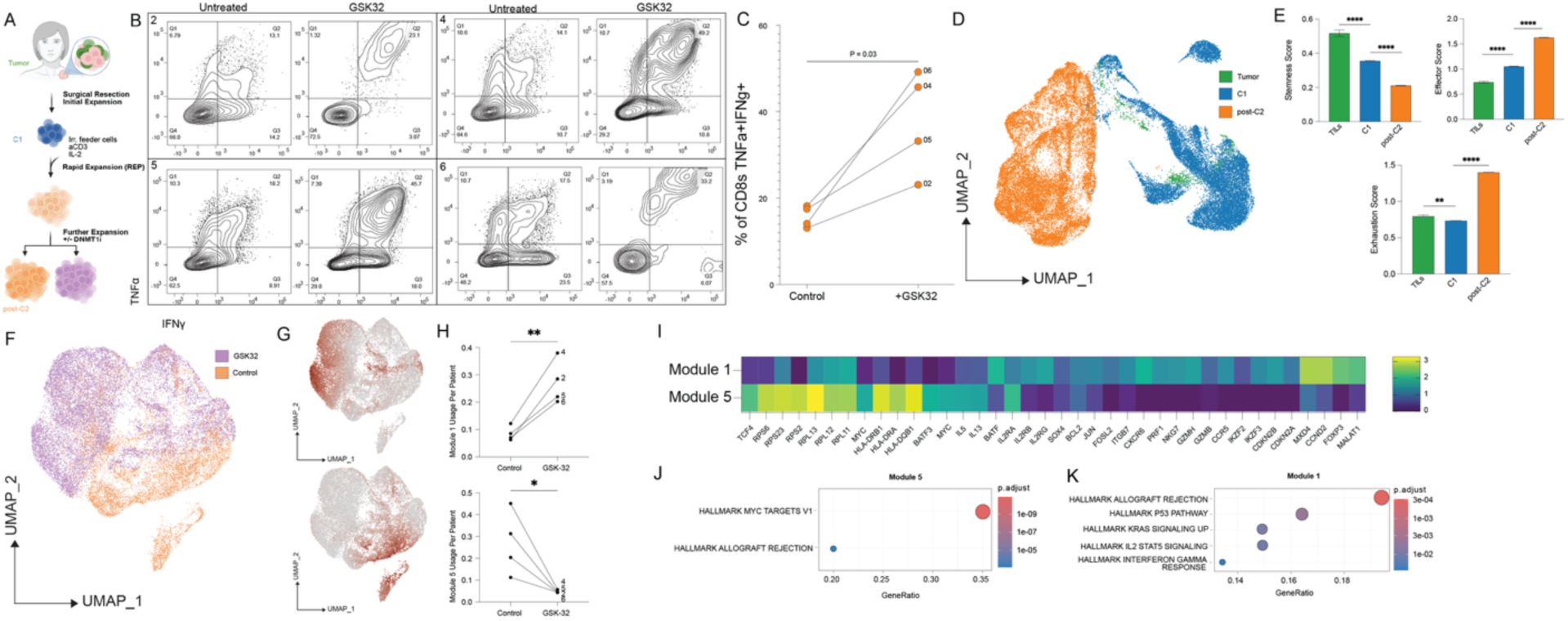
DNMT1i reinvigorates REP-TILs from patients with metastatic melanoma. (A) Schematic showing timepoints for TIL collection and DNMT1i treatment. (B) TNFα and IFNγ production by REP TILs with or without DNMT1i for patients 2,4,5 and 6. (C) Quantification of TNFα+IFNγ+ CD8⁺ T cells in the absence or presence of DNMT1i (n=4, paired t test). (D) UMAP reduction of TILs from all patients (1–6) colored by timepoint (tumor, C1, and post-C2). (E) Stemness, effector and exhaustion modules were compared between timepoints (two-sided Wilcoxon rank-sum tests applied to single-cell module scores). (F) UMAP reduction of *CD8+* TILs from post-C2 only colored by treatment (control: orange, GSK32: lilac). (G) UMAP reduction of *CD8+* TILs from post-C2 only colored by module 1 usage (top) and module 5 usage (bottom). (H) Quantification of module 1 usage (top, two-sided paired t test, *p* = 0.0097) and module 5 usage (bottom, two-sided paired t test, *p* = 0.0312) across treatment (n=4). (I) Heatmap showing gene loadings for modules 1 and 5 from the top 150 ranked gene list for each module. (J-K) ORA results for top 200 ranked genes from module 5 (J) and module 1 (K). scRNA-seq includes all patients 1–6 (tumor/C1/post-C2) while functional/CD8 analyses use the four CD8-dominant samples.

We performed single cell RNA sequencing (scRNAseq) on DNMT1i treated and untreated post-C2 TILs from patients 1-6 as well as paired samples from each patient from the original tumor sample and C1 without treatment (Fig 6A). We recovered 21,339 CD8⁺ T cells from post-C2, 20,588 from C1, and 785 from original tumor samples (Fig 6D). To determine how *ex vivo* expansion impacts TIL transcriptional state, we defined stemness (*TCF7, LEF1, IL7R, CCR7, KLF2*), effector (*GZMB, PRF1, TBX21, IFNG, TNF*), and exhaustion (*HAVCR2, LAG3, ENTPD1, CTLA4*) scores (Fig 6E). Untreated TILs progressively lost transcriptional features of stemness and gain an effector signature over time (Fig 6E, S6B). From tumor to C1, TILs transiently lost exhaustion features, but these were regained post-C2 (Fig 6E).

Next, we sought to characterize the response of the terminally exhausted post-C2 TILs to DNMT1i. Notably, DNMT1i treatment led to depletion of *CD4*⁺ TILs, but not *CD8*⁺ TILs, in post-C2, suggesting CD4⁺ TILs may not be amenable to DNMT1-mediated reprogramming (Fig S6C,D). To analyze CD8-specific responses, we re-clustered *CD8*⁺ TILs from the four CD8-dominant samples (2, 4, 5 and 6) (Fig 6F). Since clustering largely identified patient-specific clusters, we used cNMF to learn unbiased gene modules active across multiple clusters. cNMF analysis identified 7 modules, among these modules 1 scores showed positive correlation whereas module 5 scores showed negative correlation with DNMT1i across patients (p = 0.0097 and p = 0.0312, respectively, Fig 6F-H). We performed overrepresentation analysis of top genes associated with each module. Module 1 was strongly loaded for genes associated with allograft rejection (*GZMB, IL2RA, IL2RG, PRF1*) while module 5 was strongly loaded for Myc target genes (*LDHA, EEF1B2, EIF4A1, RPL22, RPS 2/3/5/6*) (Fig6I-K). Module 1 was further strongly loaded for activation-associated *JUN* and *FOSL2* and effector-associated transcription factors *BATF, IKZF2* and *IKZF3* (Fig 6). In addition to Myc targets, module 5 showed strong loadings for *MYC* itself, *IL5* and *IL13* and memory-associated transcription factor *BATF3* (Fig 6I). DNMT1i thus induced divergent programs in post-C2 CD8 TILs. Expanding, untreated CD8⁺ T cells preferentially used a program associated with Myc signaling, biosynthetic activity, and type 2 cytokine expression while DNMT1i-treated CD8⁺ T cells preferentially used a program associated with effector function, activation, and cytotoxicity.

## Discussion

The differentiation of CD8⁺ T cells proceeds along a continuum from naïve to effector, memory, and exhausted states, with epigenetic mechanisms encoding both durable protective immunity and dysfunctional responses under chronic stimulation. In acute infection and CAR-T cell models, a period of antigen withdrawal and cellular quiescence after priming is essential for the acquisition of memory whereas persistent stimulation diverts cells into an exhausted fate characterized by diminished recall and restricted proliferative potential (*9, 30, 34–37*). Exhausted CD8⁺ T cells display extensive chromatin remodeling relative to memory cells, and prior work has shown that neither antigen withdrawal nor immune checkpoint blockade fully restores a memory-like epigenetic landscape once exhaustion is established (*30, 38, 39*). These observations have led to the view that exhausted CD8⁺ T cells are epigenetically “scarred” and largely fixed in their differentiation trajectory(*30*).

We investigate whether interfering with maintenance DNA methylation can alter the trajectory of CD8⁺ T cell differentiation and enhance anti-tumor immunity. We show in a melanoma model that pharmacologic hypomethylation via DNMT1i synergizes with CTLA4-directed ICI to enrich a progenitor-exhausted (T_PEX_) CD8⁺ population in the tdLN that is transcriptionally and functionally similar to human checkpoint-responsive T_PEX_. *In vivo*, combining DNMT1i with immune checkpoint blockade shifts the tdLN CD8⁺ compartment away from naïve-like precursors towards T_PEX_ without significantly altering the frequency of T_EFF EX_ in the tumor or tdLN. The latter finding may reflect differences in the lifespan or trafficking equilibrium of terminal effectors. These data support a model that DNMT1i biases early fate choices, favoring the generation and maintenance of a resource T_PEX_ pool that can fuel effective effector responses under checkpoint blockade.

Our *in vitro* model further demonstrates that DNMT1i rescues the function of chronically stimulated CD8⁺ T cells by inducing a divergent transcriptional and regulatory state rather than returning cells to a canonical memory-like fate. Chronic PKC-dependent signaling drives a typical exhausted phenotype with diminished TNFα/IFNγ production and cytotoxicity despite increased proliferation and robust expression of exhaustion- and effector-associated genes. DNMT1i during this chronic stimulation preserves dual TNFα⁺IFNγ⁺ producers and restores tumor killing, even when drug exposure was either limited to the first 36 hours after priming or after chronic stimulation. These findings indicate that DNMT1 can act both early during repeated stimulation and after chronic stimulation signals have already been integrated to imprint exhaustion-associated programs and that transient interference with DNMT1 activity is sufficient to protect effector potential.

Transcriptionally, DNMT1i redirects exhausted CD8⁺ T cells away from activation- and terminal effector-associated programs towards stemness- and early effector-like modules. Upon restimulation, DNMT1-inhibited cells maintain higher usage of an early effector module and reduce engagement of a terminal exhaustion module. Importantly, comparison with transcriptomes of DNMT1 KO human T cells shows substantial overlap in differentially expressed genes, supporting the pharmacologic effects we observe as predominantly on-target for DNMT1 and are largely shared between mouse and human.

Unexpectedly, the large transcriptional shift that we observe arises in the context of modest and highly focal changes in the chromatin accessibility. Chronic stimulation broadly remodels chromatin relative to T_M_, with T_EX_ acquiring a globally more accessible chromatin landscape. Paradoxically, DNMT1i primarily reduces global chromatin accessibility and alters a limited subset of peaks, with substantial overlap between regions closed in DNMT1i-treated exhausted CD8⁺ T cells and memory-like CD8⁺ T cells. Upon restimulation, however, the chromatin landscape of DNMT1i-treated exhausted CD8⁺ T cells shifts toward a more memory-like state. Thus, DNMT1i does not globally “open” chromatin but rather counteracts the global increase in accessibility associated with exhaustion and likely primes a subset of regulatory regions and targets whose accessibility may be dynamically reorganized upon subsequent TCR engagement.

These findings have two implications. First, they argue that single timepoint chromatin accessibility profiles under homeostatic conditions are insufficient to classify exhausted cells as irreversibly “scarred.” Alternative epigenetic patterning that eludes the resolution of global chromatin accessibility may be differentially remodeled upon restimulation. Second, they suggest that DNMT1i primarily modulates epigenetic plasticity i.e. the capacity of chromatin to undergo state-dependent remodeling in response to extrinsic cues, rather than imposing a stable alternative, context-independent landscape. These two mechanistic proposals rely on the assumption that pharmacologic DNMT1i hypomethylates CpG sites, which then alter subsequent context-dependent epigenetic patterning. Indeed, DNA methylation analysis confirms that DNMT1i globally hypomethylates exhausted CD8⁺ T cells even in the absence of associated chromatin accessibility changes. The global hypomethylating activity of DNMT1i is expected as DNMT1 maintains methylation marks across divisions and its inhibition induces passive demethylation. The finding that DNMT1i counterintuitively leads to hypermethylation of select promoters of key genes, including *Myc* and *Atf4*, requires further investigation to evaluate DNMT1i-induced mechanisms of selective hypermethylation.

Methylation analysis as well as motif and regulon analysis highlight specific TF networks through which DNMT1i reshapes exhausted CD8⁺ T cell states. Notably, DNMT1i does not restore TCF7-family activity, consistent with prior work identifying DNMT3A rather than DNMT1 as the primary regulator of *Tcf7* in CD8⁺ T cells(*10*). This dissociation between functional rescue and TCF7-family activity aligns with the growing recognition that TCF1 expression is an imperfect surrogate for effector competence, and that progenitor-marker abundance and anti-tumor function can be uncoupled. Our data indicate that DNMT1i improves effector output through a TCF7-independent regulatory network, arguing that functional readouts, rather than TCF1⁺ frequency alone, are the appropriate metric for epigenetic reprogramming strategies. DNMT1i sustains or enhances regulatory activity of KLF, LEF1, FOX, and IKZF1 families and attenuated MYC and ATF family activity. Integrated expression and methylation analysis validates *Foxo1, Lef1, Ikzf1,* and several *Klf* members as key targets of DNMT1i-driven hypomethylation and both *Myc* and *Atf4* as stably hypermethylated and downregulated downstream of DNMT1i. This pattern suggests that DNMT1 promotes a Myc-driven biosynthetic program under chronic stimulation whereas DNMT1i enables a more quiescent, survival-competent and interferon-responsive regulatory network that permits strong effector responses when appropriately stimulated. Considering the established roles of FoxO1 and BACH2 in sustaining CD8⁺ T cell survival and stemness in chronic antigen exposure, and their recently validated function in maintaining CAR-T cell fitness, it is notable that DNMT1i coordinately hypomethylates and upregulates the Foxo1 and Bach2 regulons and enforces the Klf2/Sell (CD62L) program associated with stem-like, differentiation-competent progenitors(*40–42*). This suggests DNMT1i re-engages a maintenance module relevant for durable adoptive-cell-therapy products (*43*). The context-dependent nature of Myc in CD8⁺ T cell differentiation could explain why DNMT1i-treated cells exhibit improved cytokine production and cytotoxicity despite reduced proliferation and Myc activity under chronic stimulation, limiting Myc-driven biosynthetic pressure may protect cells from terminal dysfunction and allow early effector programs to be deployed (*44, 45*).

Rather than reverting exhausted CD8⁺ T cells to a canonical memory state, DNMT1i re-licenses discrete survival- and memory-associated programs, exemplified at the *Il7r* locus. There, DNMT1i drove promoter-proximal and intragenic hypomethylation, restimulation-dependent chromatin re-opening, and transcriptional de-repression of *Il7r*, and potentiated IL-7–induced STAT5 activity (Fig. 5). Notably, this re-engagement of the IL-7R–STAT5 axis occurred without restoration of TCF7-family activity, consistent with DNMT3A rather than DNMT1 governing *Tcf7* (*10*). DNMT1 and DNMT3A may thus divide control of the memory program, with DNMT1 restraining *Il7r* and other maintenance-methylated survival loci while DNMT3A patterns *Tcf7*, such that DNMT1 inhibition restores IL-7 responsiveness without fully engaging the TCF7 program and yields the non-canonical, "hybrid" state we observe rather than a complete memory reset. That DNMT1i improves function without re-establishing the TCF7 program is in keeping with recent evidence that restoring TCF1 is itself insufficient to reverse exhaustion: in chronic viral infection, conditional TCF1 over-expression in committed intermediate-exhausted CD8⁺ T cells produces only mild transcriptional and epigenetic changes and fails to regenerate the stem-like progenitor pool, likely because TCF1 cannot remodel an already-fixed chromatin landscape (*46*). Our data suggest that intervening upstream, at the maintenance-methylation machinery that constrains this landscape, rather than at a single downstream transcription factor, may offer an alternative route to restoring effector competence in chronically stimulated CD8⁺ T cells. Because IL-7R marks anti-tumor memory CD8⁺ T cells with superior efficacy (4), restoration of IL-7R–STAT5 competence provides a plausible mechanistic link between DNMT1i and the restored effector function we observe, including in patient-derived TILs. We note that STAT5 reporter activity was measured in a heterologous 293T system; direct confirmation that DNMT1i amplifies IL-7–STAT5 signaling in primary exhausted CD8⁺ T cells will further substantiate this axis.

Finally, DNMT1i improves effector function in highly differentiated, clinically relevant human TIL products from patients with metastatic melanoma who had progressed on prior ICI. In extensively expanded TILs, DNMT1i increases the fraction of TNFα⁺IFNγ⁺ CD8⁺ T cells and induced gene modules enriched for cytotoxicity, activation, and allograft rejection signatures, while untreated CD8⁺ TILs preferentially engaged a Myc-driven, biosynthetic, and type 2 cytokine-associated program. These results suggest that DNMT1i can rebalance transcriptional programs in terminally expanded human CD8⁺ TILs, enhancing functional quality even at late stages of ex vivo manipulation. However, further studies are required to fully understand the functional implications of the observed phenotypic switch. Notably, as DNMT1i preferentially drove activity of an activation and cytotoxicity program over a Myc-driven biosynthetic program in TILs, the transcriptional consequences of DNMT1i may be cell state-dependent and require further investigation at the chromatin and methylation level and interrogation in TILs from different differentiation timepoints. Together, this supports a model in which DNMT1 regulates how CD8⁺ T cells integrate chronic stimulation, biosynthetic stress, and differentiation cues into durable epigenetic and functional states.

Collectively, our data suggest that DNMT1 does not act as a master “on/off” switch for exhaustion but rather as a modifier of fate branching and epigenetic plasticity under chronic stimulation. DNMT1i in this context does not simply induce a canonical memory state; rather, it establishes a distinct, drug-conditioned trajectory in which exhausted CD8⁺ T cells maintain superior effector capacity while remaining transcriptionally and epigenetically distinct from both T_EX_ and T_M_. Importantly, our data do not indicate reversion of terminally differentiated cells to a bona fide progenitor or memory state, consistent with the field’s consensus that terminal exhaustion is not reversed by current interventions. Rather, DNMT1i acts on fate branching and plasticity to preserve effector competence within a distinct, drug-conditioned state. Given the multiplicity of epigenetic enzymes that shape CD8⁺ T cell differentiation, targeting any single enzyme is likely to yield such “hybrid” states rather than fully reset differentiation towards an alternative physiologic trajectory. Systematic perturbation of individual epigenetic regulators, combined with longitudinal function and multi-omic profiling, will be necessary to map the relative contributions of each pathway and to rationally design immune-epigenetic therapeutic strategies that enhance durable anti-tumor immunity without compromising lineage identity nor long-term persistence.

### Study Design

The objective of this study was to test whether DNMT1 inhibition alters CD8⁺ T cell differentiation and function during chronic stimulation and in the antitumor immune response. To address this, we combined in vivo mouse melanoma experiments, in vitro CD8⁺ T cell exhaustion assays, bulk transcriptomic and epigenomic profiling, and single-cell RNA-sequencing of mouse and human tumor-infiltrating lymphocytes. Mouse experiments evaluated tumor growth, survival, and CD8⁺ T cell states following treatment with DNMT1 inhibitor, anti-CTLA-4, or the combination. Single-cell RNA-sequencing of tumors and tumor-draining lymph nodes was used to quantify CD8⁺ T cell states in a using mice as replicates. In vitro experiments used independently generated CD8⁺ T cell cultures as biological replicates for functional assays, bulk RNA-seq, and ATAC-seq. Human TIL experiments analyzed paired treated and untreated samples from the same patients. Mouse tumor experiments were performed in two independent cohorts. No formal randomization was used, and investigators were not blinded. Mice that died from causes unrelated to tumor burden were excluded from survival analyses. For single-cell experiments, low-quality cells and doublets were excluded using standard quality-control criteria described in Methods. No statistical methods were used to predetermine sample size. Details of biological replicates and group sizes are provided in the figure legends.

### Statistical Analysis

Statistical analyses were performed using GraphPad Prism 10 and R. A two-sided P value < 0.05 was considered statistically significant. Throughout the figures, *P < 0.05, **P < 0.01, and ***P < 0.001. Mouse survival was analyzed using Kaplan–Meier curves with log-rank tests. Flow cytometry–based cytokine assays and in vitro tumor-killing assays were analyzed using two-tailed unpaired t tests, with independent cultures treated as biological replicates. For human single-cell gene-program analysis, module usage values were compared between paired treated and untreated samples from the same patient using paired two-tailed t tests. For bulk RNA-sequencing, differential expression was assessed using DESeq2 with Benjamini–Hochberg correction for multiple testing. For ATAC-sequencing, differential chromatin accessibility was assessed using DiffBind with DESeq2 normalization and FDR-based significance thresholds. For single-cell RNA-sequencing, differential expression between clusters was performed using Seurat’s Wilcoxon rank-sum test, and pseudobulk comparisons were performed using DESeq2 with individual mice as replicates. Gene set enrichment analyses used Benjamini–Hochberg correction. No outliers were excluded from any analysis.

### Mice

Animal care and experimental procedures were conducted in accordance with protocols approved by the Institutional Animal Care and Use Committee (IACUC) of Yale University. C57BL/6J (B6) mice were purchased from The Jackson Laboratory (stock no. 000664) and bred in house. dsRED-expressing TCR transgenic P14 mice were obtained from the laboratory of Nikhil Joshi at Yale University. Mice were housed (up to five per cage) in a controlled facility with a 12-hour light/dark cycle, ambient temperature of approximately 23 °C, and ad libitum access to food, water, and enrichment nestlets. Both male and female mice aged 6 to 12 weeks were used for in vitro experiments and were age- and sex-matched across experimental groups. No special diets or water supplementation were used. Euthanasia was performed by isoflurane overdose followed by cervical dislocation in accordance with institutional guidelines. Investigators were not blinded to group allocation during experiments.

### Mouse tumor injections

The YUMM1.7 melanoma cell line expressing the LCMV gp33–41 epitope (YUMM1.7-gp33) was used for all in vivo tumor experiments. Cells were maintained in culture under standard conditions and routinely tested for mycoplasma contamination. For tumor implantation, 7-week-old C57BL/6J mice were injected subcutaneously in the right flank with YUMM1.7-gp33 cells suspended in sterile phosphate-buffered saline. Tumors were allowed to establish for 7 days prior to initiation of treatment. Mice were then treated every other day for a total of five doses with vehicle, DNMT1 inhibitor, anti-CTLA-4, or combination therapy as indicated. DNMT1 inhibitors and anti-CTLA-4 antibody were administered intraperitoneally using separate syringes and needles for each compound. Anti-CTLA-4 was administered at a dose of 25 μg per injection. DNMT1i was administered at a dose of 85 μg per injection. Tumor size was monitored at regular intervals, and mice were euthanized when tumors reached predefined humane endpoints in accordance with institutional guidelines. All treatments and measurements were performed in accordance with Yale IACUC-approved protocols.

### Mouse single cell CITE-seq

Tumors and tumor-draining lymph nodes (TDLN) were harvested from mice on day 12 post tumor injection. Three mice per treatment group were included, and samples were multiplexed using sample-specific hashtags and antibody-derived tags (CITE-seq). Single-cell suspensions were prepared from tumors and tdLNs and processed for single-cell RNA-sequencing with paired antibody-derived tag (ADT) and TCR V(D)J profiling. Libraries were generated using the 10x Genomics Chromium platform following the manufacturer’s protocol for 5′ gene expression, CITE-seq, and V(D)J enrichment. Raw base call files were demultiplexed and processed using Cell Ranger Multi to generate gene expression, antibody-derived tag, and TCR contig matrices for each sample. Raw feature-barcode matrices were loaded for each library into Seurat. Cells were filtered based on quality control thresholds for number of detected genes, total UMI counts, and mitochondrial gene content. Gene expression data were log-normalized, highly variable genes were identified using the vst method, and expression values were scaled. Principal component analysis was performed on variable genes, and nearest-neighbor graphs were constructed for clustering and visualization. Cells were embedded using UMAP based on principal components. Clusters were identified using a graph-based Louvain algorithm and manually annotated on the basis of canonical marker genes. To perform replicate-aware comparisons, cells were grouped by mouse, condition, and cluster identity and aggregated using Seurat’s AggregateExpression function to generate pseudobulk expression profiles. These pseudobulk profiles were used to compare gene expression between conditions within defined CD8⁺ T-cell subsets using DESeq2-based statistical testing. To model differentiation trajectories among CD8⁺ T-cell states, Seurat objects were converted to Monocle3 cell data sets. Data were preprocessed, embedded in UMAP space, and clustered, and a principal graph was learned to infer developmental trajectories. Cells were ordered along pseudotime, and genes varying along the trajectory were identified using graph-based differential expression testing. To compare mouse CD8⁺ T-cell states with human melanoma CD8⁺ T-cell populations, one-to-one human–mouse orthologs were obtained from Ensembl. Average gene expression per cluster was computed separately for mouse and human datasets, and Pearson correlation was calculated between ortholog-matched expression matrices. Correlation matrices were visualized to quantify similarity between mouse and human CD8⁺ T-cell states.

### In vitro CD8 exhaustion model

Naïve CD8⁺ T cells were isolated from spleens and peripheral lymph nodes (inguinal, brachial, and axillary) of 7–8-week-old P14 TCR-transgenic mice using magnetic negative selection (Miltenyi). In parallel, dendritic cells (DCs) were isolated from spleens of age-matched C57BL/6J mice. Splenocytes were pulsed for 4 h at 37 °C with gp33 peptide (5 μg/mL) and poly(I:C) (20 μg/mL), after which pan-DCs were enriched by magnetic isolation (Miltenyi). Purified P14 CD8⁺ T cells were co-cultured with peptide-pulsed DCs at a 10:1 CD8:DC ratio at a density of 1×10⁶ cells/mL in T-cell medium (RPMI supplemented with 10% FBS, non-essential amino acids, penicillin/streptomycin, 20 mM HEPES, 1 mM sodium pyruvate, and 35 ng/mL recombinant human IL-2). To induce chronic stimulation (“exhaustion”), cultures were split into +PMA and −PMA groups immediately following priming. For +PMA conditions, cells were exposed to PMA (1 μg/mL) for 5 min at 37 °C, washed, and then cultured long-term in the presence of low-dose PMA (20 ng/mL). PMA was replenished every 48 h during media changes. Control cultures were treated with vehicle. DNMT1 inhibitors (RG108 or GSK32) and IL-2 were replenished every 48 h. On day 7, cells from each condition were split into restimulated and non-restimulated groups. Restimulation was performed with gp33 peptide (5 μg/mL) in the presence of GolgiPlug for 4 h at 37 °C; control wells received vehicle with GolgiPlug. Cells were then stained for surface markers (CD45, CD3, CD8, CD4) followed by intracellular staining for TNF-α and IFN-γ using the FoxP3 intracellular staining kit (eBioscience) and analyzed by flow cytometry. To assess cytotoxic function, gp33-expressing YUMM melanoma cells (YUMM-G1-gp33-GFP) were plated at 10,000 cells per well in 96-well flat-bottom plates in T-cell medium supplemented with IFN-γ (100 ng/mL) 24 h prior to co-culture. On day 7, live CD8⁺ T cells were enriched using a dead-cell removal kit (Miltenyi) and added to tumor cells at a 1:10 T-cell:tumor ratio. Co-cultures were incubated for 24 h in IL-2–containing medium. Cells were then harvested, stained for viability and T-cell markers, and tumor cell killing was quantified by flow cytometry. For spectral flow cytometry experiments, cells were stained for viability and T-cell markers and data was acquired on a Cytek Aurora. FlowJo was used for analysis and the PhenoGraph algorithm was used for unsupervised clustering (K-means=120).

### Bulk RNA Sequencing of P14 CD8 T Cells

CD8⁺ T cells were harvested from in vitro cultures under the indicated stimulation and drug conditions in triplicates. Total RNA was extracted using the RNeasy kit (Qiagen). RNA quality and concentration were assessed prior to library preparation. Poly(A)-selected RNA-seq libraries were prepared and sequenced by the Yale Center for Genome Analysis (YCGA). Sequencing was performed with paired-end reads to sufficient depth for differential expression analysis. Transcript-level abundances were quantified using kallisto and differential expression analysis was performed using DESeq2. Genes with low counts were filtered prior to analysis (min. 5 reads in at least 2 samples). DE testing was carried out using the Wald test. LFC shrinkage was performed using the ashr method. Genes were considered differentially expressed based on adjusted p values (Benjamini–Hochberg correction) and effect size thresholds as specified in figure legends. Gene set enrichment analysis was performed on differential expression results using the clusterProfiler package with gene sets derived from MSigDB, including Hallmark (H), curated (C2), and immunologic (C7) collections. To identify latent transcriptional programs across conditions, consensus non-negative matrix factorization (cNMF) was applied to normalized gene expression data. Normalized DESeq2 counts were used as input after selecting the top 2,000 most variable genes across samples. Ribosomal, mitochondrial, and housekeeping genes were excluded prior to factorization. The optimal number of modules was determined by rank estimation across ranks 2–8 using repeated runs of the Brunet algorithm. A four-module solution was selected based on stability and interpretability, and NMF was run with 50 iterations to obtain robust gene-by-module and sample-by-module matrices. Module usage across conditions was calculated from sample-by-module coefficients and visualized to compare pathway activity across stimulation and drug conditions. To compare DNMT1i treatment of P14 CD8⁺ T cells to DNMT1KO in human T cells, differentially expressed genes (adjusted *p* < 0.05) from the drug-treated dataset and the published knockout dataset from Li et al. were compared. To formally assess whether differential expression in the drug-treated dataset was enriched among genes differentially expressed in the knockout dataset, analysis was restricted to the set of genes shared between both datasets. Each gene was classified as differentially expressed or not (*padj* < 0.05). A contingency table was constructed comparing differential expression status across datasets, and statistical significance of the overlap was assessed using a chi-squared test of independence.

### Bulk ATAC Sequencing of P14 CD8 T Cells

ATAC-seq libraries were generated from in vitro CD8⁺ T-cell samples as described above using the Active Motif ATAC-Seq Kit (53150). Libraries were sequenced on Illumina platforms. Raw sequencing data were processed using the ENCODE ATAC-seq pipeline, which performs adapter trimming, alignment to the mouse genome (mm10), filtering of low-quality and duplicate reads, mitochondrial read removal, Tn5 shifting, and generation of normalized signal tracks and peak calls. Peaks were called on individual samples, and a union peak set across all samples was constructed for downstream comparative analyses. Read counts for each peak in the union set were quantified for all samples and used for differential accessibility testing with DiffBind, using DESeq2 for normalization and statistical modeling. Pairwise contrasts were defined to compare TM, TEX, TEX*, and TEX+D conditions. Differentially accessible regions (DARs) were identified using an FDR threshold of 0.05, and log2 fold-changes were used to determine relative accessibility between conditions. To link differential chromatin accessibility to genes, DARs were annotated to nearby transcription start sites using ChIPseeker and the TxDb.Mmusculus.UCSC.mm10.knownGene annotation. For transcription factor footprinting, we applied GimmeMotifs (*47*). To resolve transcription factor binding at base-pair resolution, we applied ChromBPNet, a deep-learning framework that models Tn5 bias and learns sequence-specific footprints directly from ATAC-seq data (*31*). ChromBPNet models were trained on each condition to predict chromatin accessibility and infer transcription factor binding events, enabling accurate identification of motif-specific occupancy independent of Tn5 insertion bias. Using the learned motif profiles, we quantified motif footprint density in the vicinity of gene promoters and regulatory regions across TM, TEX, TEX*, and TEX+D conditions. To infer transcription factor (TF) activity from chromatin accessibility, we used decoupler with the CollecTRI regulon database, which links TFs to their experimentally supported target genes. TF activity scores were computed from accessibility-weighted motif and target gene signals.

### Patient-derived TIL expansion and single-cell RNA sequencing

Treated and untreated tumor-infiltrating lymphocytes (TILs) were harvested at the indicated time points for functional and transcriptional profiling. For functional assessment, continuously stimulated TIL cultures (anti-CD3/anti-CD28 with IL-2) were restimulated with phorbol 12-myristate 13-acetate (PMA) and ionomycin using Cell Stimulation Cocktail in the presence of brefeldin A. Cells were then stained for viability, CD45, CD4, and CD8, followed by intracellular staining for TNFα and IFN-γ and analyzed by flow cytometry. For single-cell RNA-sequencing, TILs were processed using the 10x Genomics Chromium platform with paired gene expression, antibody-derived tag (CITE-seq), and TCR V(D)J libraries. Cells from multiple patients and treatment conditions were multiplexed using hashtag antibodies. Libraries were sequenced and processed using Cell Ranger to generate gene expression, ADT, and TCR contig matrices. Gene expression and ADT matrices were loaded into Seurat, and cells were filtered based on gene number and mitochondrial transcript content. RNA data were log-normalized, variable genes were identified, and data were scaled prior to principal component analysis.

Graph-based clustering was performed, and UMAP was used for visualization. ADT data were normalized using centered log-ratio normalization and used to confirm surface marker expression. Hashtag demultiplexing was performed using ADT counts. TCR clonotype was processed and appeneded using scRepertoire. Cell clusters were annotated based on canonical marker genes. To analyze differentiation trajectories, Seurat objects were converted to Monocle3 cell data sets. Data were preprocessed, embedded in UMAP space, and principal graphs were learned to model lineage relationships. Cells were ordered in pseudotime, and gene expression dynamics were examined along inferred trajectories. For visualization and scoring, curated gene sets representing stem-like, effector, and exhausted T-cell states were used to compute module scores in Seurat using AddModuleScore. Scores were compared across time points and treatment conditions using Wilcoxon rank-sum tests. For gene program analysis, non-negative matrix factorization (cNMF) was applied to CD8⁺ T cells. Module usage values and gene loading matrices were computed, and module-specific gene sets were extracted from the fitted model. For each module, genes were ranked by their contribution to module loadings, and top-ranking genes were used for downstream analyses. Module-specific gene sets were analyzed using over-representation analysis against MSigDB Hallmark gene sets. Enrichment was performed using clusterProfiler with Benjamini–Hochberg correction. To test whether DNMT1 inhibition altered program activity, module usage values were compared between paired treated and untreated samples from the same patient. Statistical testing was performed using paired t-test.

## Acknowledgments

GM is supported by the Melanoma Research Alliance Young Investigator Award, Yale Center for Clinical Investigation, Dermatology Foundation, National Cancer Institute and the Yale SPORE in Skin Cancer (P50CA121974-18). MWB is supported by the Melanoma Research Alliance, NCI (U01CA238728-05), and Department of Defense. MGD is supported by the T32 in Dermatology at Yale University (2T32AR007016-50). We thank Simon F. Roy, Veronica T. Brooks, and Karine Flem-Karlsen for help with different aspects of this work.

## Author Contributions

Conceptualization, GM and MGD; data acquisition and methodology, MGD, GM, SM, KP, PG, DS, SK; data curation, MGD, GM; formal analysis, MGD, KO, GM, SM; supervision, GM, MWB, KSA, SK; visualization MGD, SM, KO; writing—original draft, MGD; writing—review and editing, MGD, GM, KSA, MWB.

## Data and Materials Availability

All sequencing data generated in this study, including bulk RNA-seq, bulk ATAC-seq, mouse single-cell RNA-seq, and human TIL single-cell RNA-seq, will be deposited in the Sequence Read Archive (SRA) upon acceptance of the manuscript.

