## Supplementary Figures for "Targeting DNMT1 augments anti-tumor CD8⁺ T cell function"

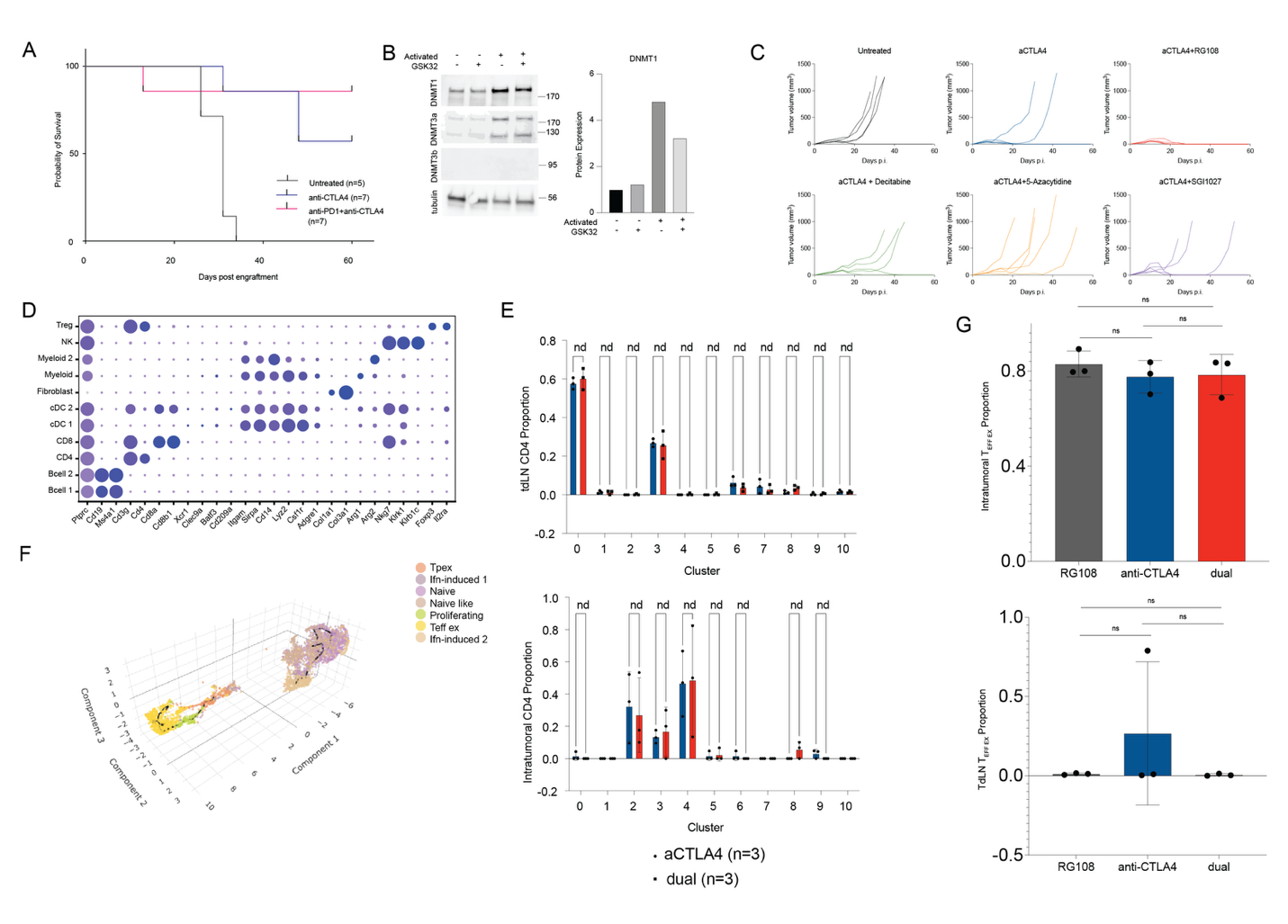

**Figure S1.** (A) Survival for untreated, aCTLA4, and aCTLA4+aPD1 treated mice. (B) Western blot of mouse naïve or activated CD8+ T cells untreated or treated with GSK32 (left) and quantification (right). (C) Tumor growth curves for treatment with (top) vehicle, aCTLA4, aCTLA4+RG108 and (bottom) aCTLA4+decitabine, aCTLA4+5-Azacytidine, aCTLA4+SG1027. (D) Dot plot showing expression of key genes defining immune cell clusters from mouse scRNAseq. (E) Bar plots of proportion of tdLN (top) and intratumoral (bottom) CD4s across clusters split by treatment. (F) 3D visualization of Monocle pseudotime trajectory. (G) Quantification of proportion of (top) intratumoral T_EFF EX_ and (bottom) tdLN T_EFF EX_ across treatment groups (two-sided unpaired t test).

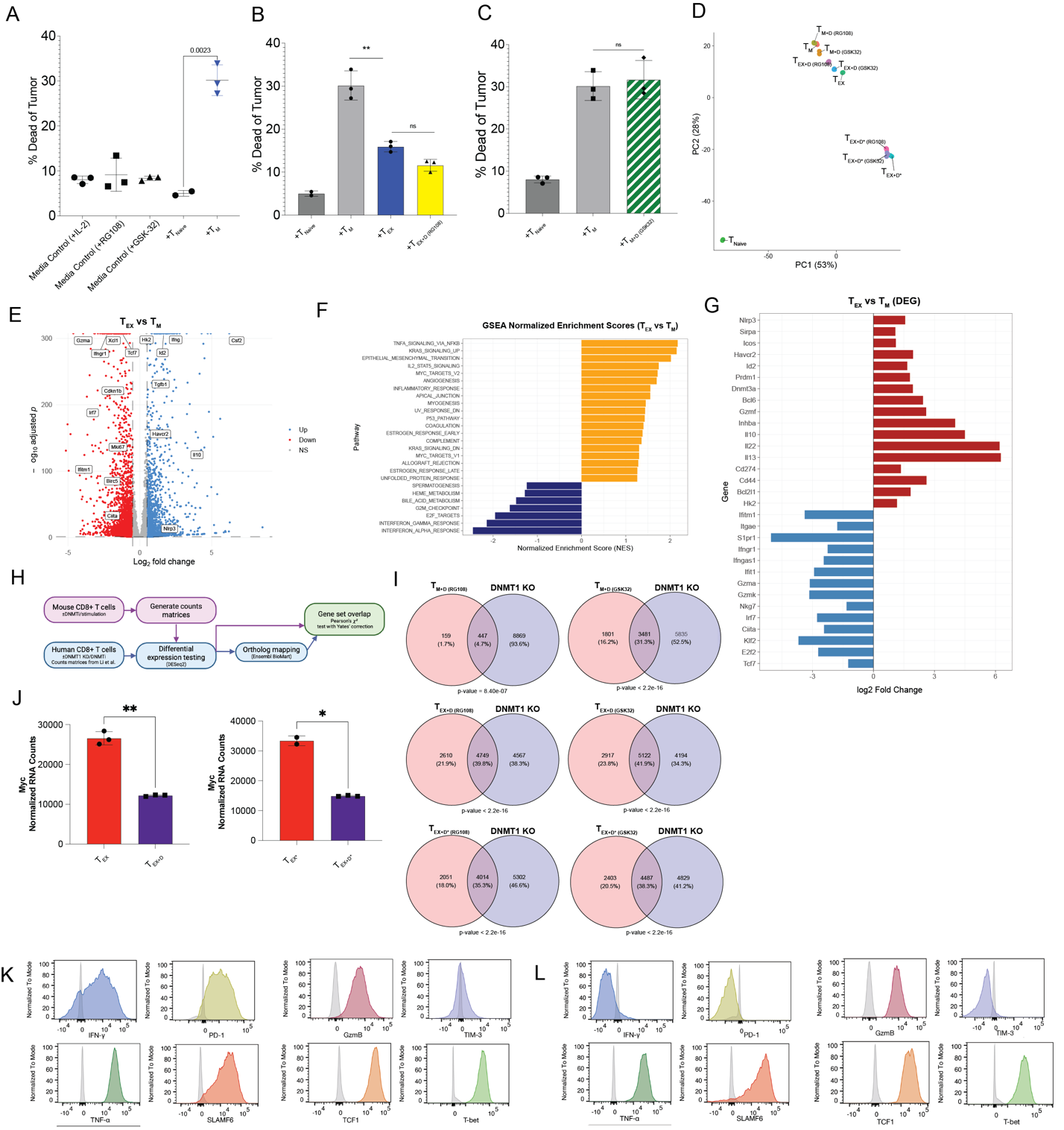

**Figure S2**. (A) Quantification of T_Naive_ vs T_M_ with a two-sided unpaired t test. (B) Quantification of T_EX_ vs T_EX+D_ with RG108 with a two-sided unpaired t test. (C) Quantification of tumor killing capacity of CD8s treated with GSK32 in the absence of chronic stimulation (T_M+D_) versus control (T_M_) with a two-sided unpaired t test. (D) PCA of bulk RNA-seq samples. (E) Volcano plot showing differentially regulated genes between T_EX_ and T_M_. Cutoff for adjusted p-value < 0.05. (F) GSEA result top terms for T_EX_ vs T_M_ (p adj < 0.05). (G) Barplot showing top DEG for T_EX_ vs T_M_ (p adj < 0.05). (H) Schematic showing workflow for comparison of DNMT1i-treated mouse CD8 DEG and DNMT1KO human T cell DEG from Li et al. (I) Venn diagrams showing overlap of DEG between labelled conditions versus respective untreated or sg control (p adj < 0.05). Overlap quantified using a chi-squared test of independence. (J) *Myc* normalized expression between conditions (n=3). (K) Spectral flow expression of indicated markers in MC4 (colored histogram) relative to mean expression across all clusters (grey). (L) Spectral flow expression of indicated markers in MC14 (colored histogram) relative to mean expression across all clusters (grey).

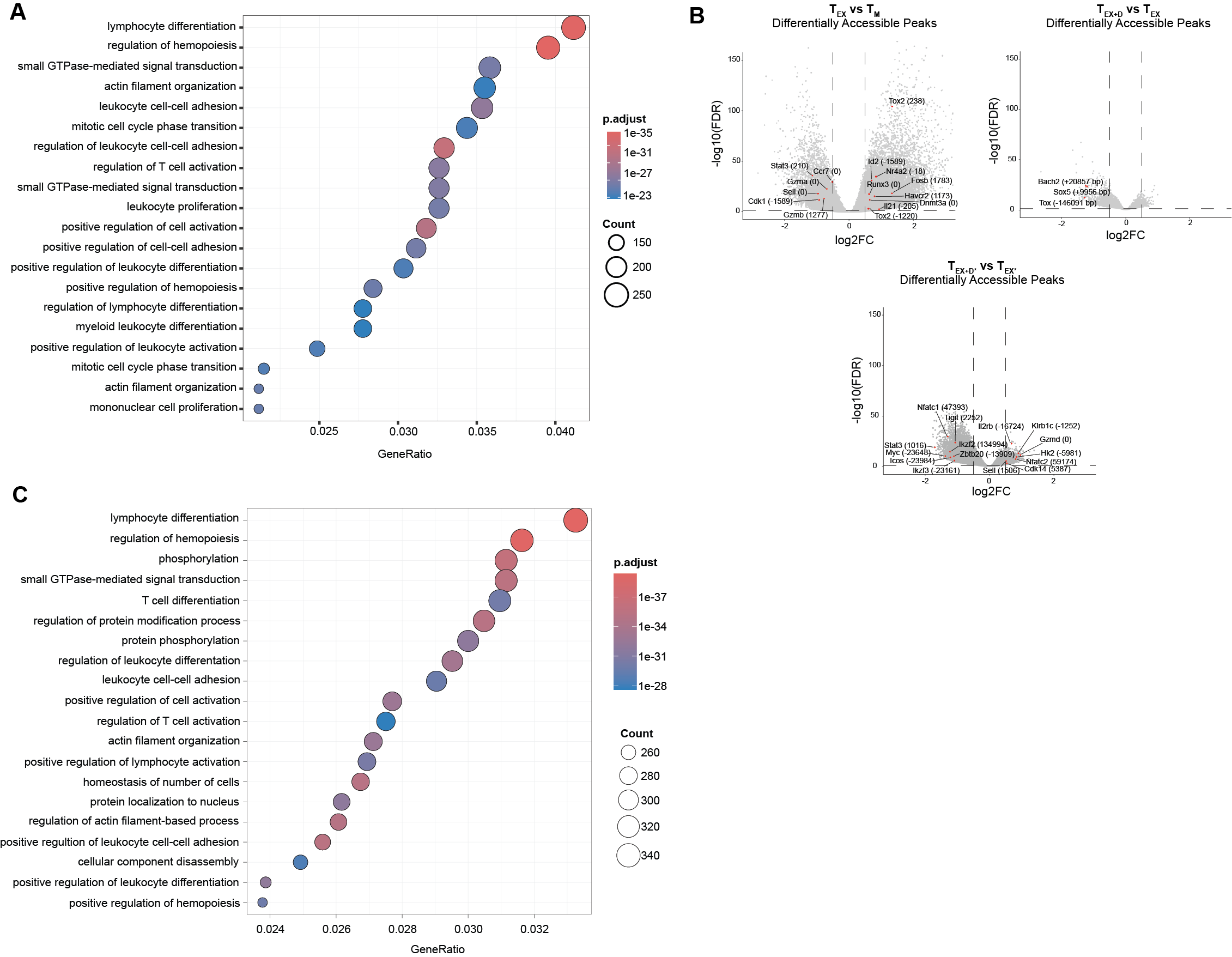

**Figure S3**. (A) ORA results for shared peaks between T_EX+D_ vs T_EX_ and T_M_ vs T_EX._ (B) Volcano plots showing differentially accessible peaks between (top left) T_EX_ vs T_M_, (top right) T_EX+D_ vs T_EX_, and (bottom) T_EX+D*_  vs T_EX*_. Adj p-value < 0.05. Selected peaks annotated mapping to immune-related gene regulatory regions with distance to TSS. (C) ORA results for shared peaks between T_EX+D*_ vs T_EX*_ and T_M*_ vs T_EX*._

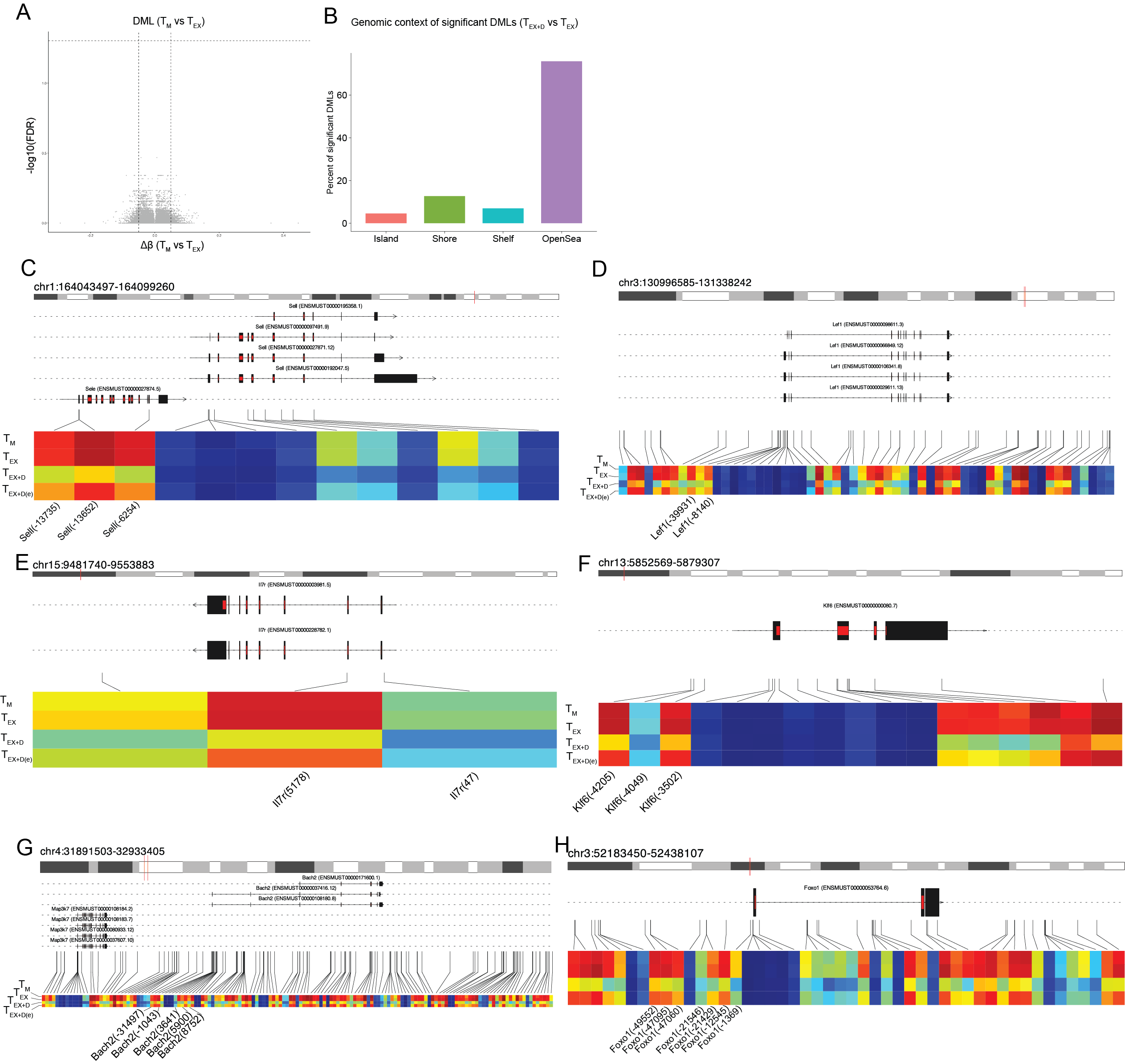

**Figure S4.** (A) Volcano plot of methylation between TEX and TM, showing no significantly differentially methylated loci. (B) Bar plot showing CpG context of differentially methylated loci (FDR < 0.05) between TEX+D and TEX. (C-H) Track-style plots showing differentially methylated loci near specific genes (*Sell, Lef1, Il7r, Klf6, Bach2, Foxo1*).

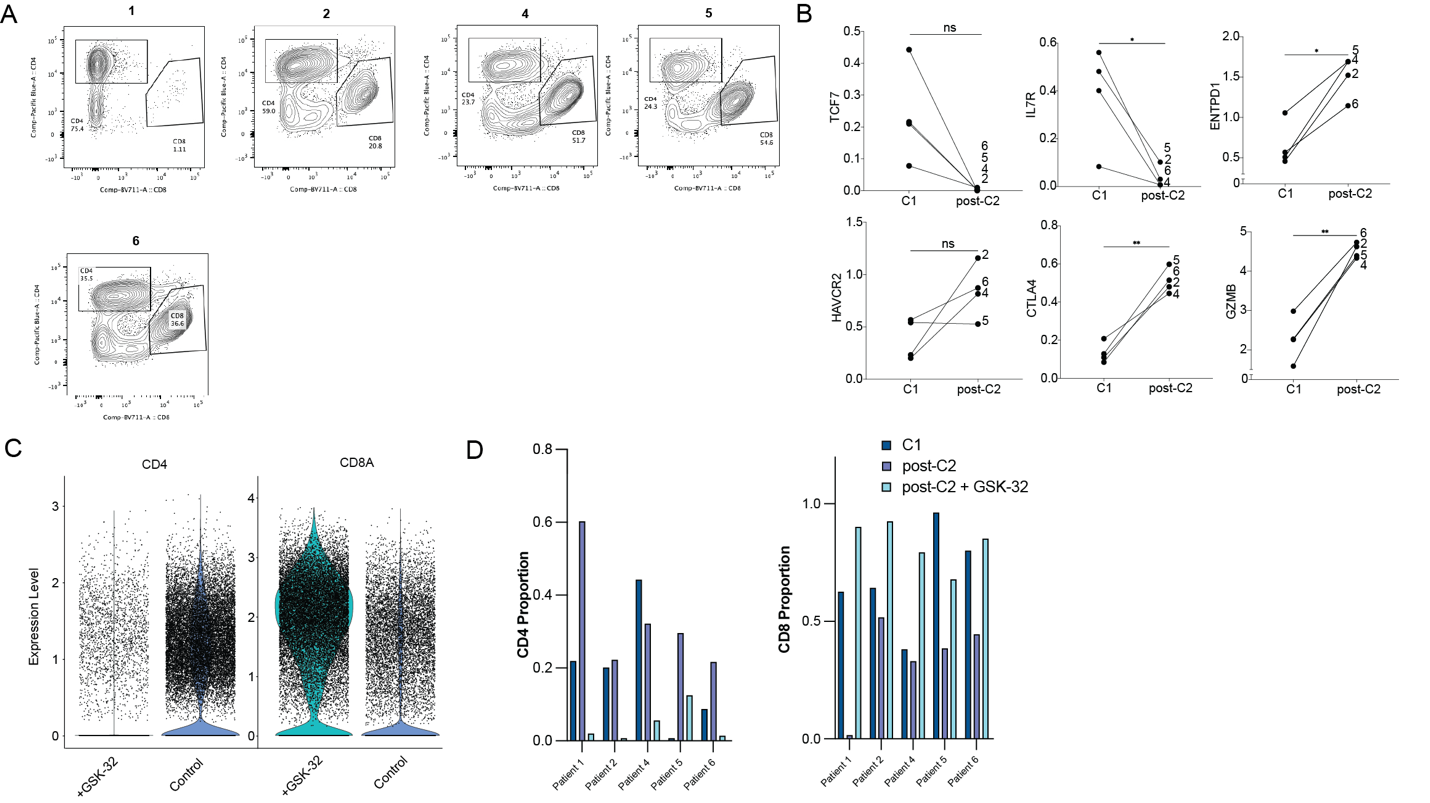

**Figure S6**. (A) Proportion of CD4+ and CD8+ TILs in post-C2 samples for patients 1-6. (B) Normalized expression of *TCF7, IL7R, ENTPD1, HAVCR2, CTLA4,* and *GZMB* in *CD8+* TILs for treated and un-treated TILs (n=4, two-tailed paired t test). (C) (left) *CD4* and (right) *CD8A* single cell expression level for untreated and treated TILs (n=5). (D) Proportion of (left) *CD4*+ and (right) *CD8+* TILs split by timepoint/treatment and patient from scRNA-seq.
